# Opposing contributions of GABAergic and glutamatergic ventral pallidal neurons to motivational behaviours

**DOI:** 10.1101/594887

**Authors:** Marcus Stephenson-Jones, Christian Bravo-Rivera, Sandra Ahrens, Alessandro Furlan, Carolina Fernandes-Henriques, Bo Li

## Abstract

The ventral pallidum (VP) is critical for invigorating reward seeking and is also involved in punishment avoidance, but how it contributes to such opposing behavioural actions remains unclear. Here we show that GABAergic and glutamatergic VP neurons selectively control behaviour in opposing motivational contexts. *In vivo* recording combined with optogenetics in mice revealed that these two populations oppositely encode positive and negative motivational value, are differentially modulated by animal’s internal state and determine the behavioural response during motivational conflict. Furthermore, GABAergic VP neurons are essential for movements towards reward in a positive motivational context, but suppress movements in an aversive context. In contrast, glutamatergic VP neurons are essential for movements to avoid a threat but suppress movements in an appetitive context. Our results indicate that GABAergic and glutamatergic VP neurons encode the drive for approach and avoidance, respectively, with the balance between their activities determining the type of motivational behaviour.

## INTRODUCTION

The decision to approach or avoid depends on the situation in the environment and the internal state of the animal. For example, thirst may encourage animals to seek a water source, but a sated animal may not find this goal worth the energy expenditure or risk (Ydenberg, 1986). Equally an extremely thirsty animal may approach a water source despite the known risk of predators.

A key region involved in goal-directed motivation is the ventral pallidum (VP). The VP is the major output structure of the ventral basal ganglia (Heimer et al., 1997). It receives projections from areas including the nucleus accumbens (NAc), prefrontal cortex and basolateral amygdala, and transmits information to multiple brain regions involved in motor control and motivation, such as the ventral tegmental area (VTA), lateral habenula (LHb), mediodorsal thalamus and pedunculopontine tegmental nucleus (Haber and Knutson, 2010). This connectivity places the VP in an ideal location to transform information about the expected value of stimuli into motivation (Mogenson et al., 1980). Indeed, a large body of work, comprehensively reviewed by others (Humphries and Prescott, 2010; Root et al., 2015; Smith et al., 2009; Stephenson-Jones, 2019), has identified the VP as a crucial driver of reward-seeking behaviour. For example, the VP is important for the normal hedonic reactions to sucrose (Farrar et al., 2008), and lesions to the VP decrease an animal’s willingness to work for reward (Farrar et al., 2008; Richard et al., 2016). Conversely, rats will work to electrically self-stimulate their VP (Panagis et al., 1995; Panagis et al., 1997), and pharmacological activation and disinhibition of the VP can both trigger feeding in sated animals (Stratford et al., 1999).

The VP is also implicated in avoidance behaviours (Stephenson-Jones, 2019; Wulff et al., 2018), as intra-VP mu-opioid activity is sufficient to drive conditioned place aversion (Skoubis and Maidment, 2003), and activating mu opioid receptors in the VP can impair conditioned taste avoidance (Inui and Shimura, 2017). In a similar manner, disinhibiting the VP through injections of the GABAergic antagonist bicuculline induces anxiety-related behaviours and increases avoidance in an approach/avoidance task in primates (Saga et al., 2017; Smith and Berridge, 2005). These findings suggest that the VP plays a role in the motivation to both seek reward and avoid punishment.

While the VP appears critical for motivating behaviour in appetitive and aversive contexts, how this structure contributes to these opposing motivational drives is unknown. One possibility is that separate populations of VP neurons drive opposing motivated behaviours. In line with this idea, *in vivo* recordings in the VP have identified two main types of neurons that are activated by the prediction of either reward or punishment (Richard et al., 2016; Saga et al., 2017; Tachibana and Hikosaka, 2012). These neurons encode information related to the expected or incentive value of stimuli (Richard et al., 2016; Tian et al., 2016; Tindell et al., 2006) and their responses are modulated by the internal state of the animal (Tindell et al., 2006). These different populations of VP neurons may be molecularly distinct, as recent optogenetic activation experiments revealed that activation of GABAergic or glutamatergic VP neurons is reinforcing or aversive, respectively (Faget et al., 2018; Knowland et al., 2017b; Tooley et al., 2018b). Altogether this body of work points to the possibility that separate populations of VP neurons play selective roles in appetitive or aversive motivation.

Here our aim was to test this hypothesis and determine if GABAergic and glutamatergic VP neurons play opposing roles in the motivation to approach or avoid. Our aim was to determine how these populations encode appetitive and aversive motivational value, examine what roles they play in appetitively or aversively motivated behaviour, and determine how their interactions influence the overall decision to approach or avoid.

## RESULTS

### Mapping functional classes onto neurochemical identities in the VP

In order to determine what GABAergic and glutamatergic VP neurons are engaged by different motivated behaviours, we trained head-fixed mice on a Pavlovian conditioning task. Each trial started with illumination of a house light and proceeded with presentation of an auditory conditioned stimulus (CS) announcing the delivery of one of the following unconditioned stimuli (USs): 0, 1 or 5 μl of water in reward blocks, and 0, 100 or 500 ms of air puff blowing to the face in punishment blocks (Figure S1A, Figure 1A, and Methods). As training progressed, mice began licking in response to the reward predicting cues. The lick rate was significantly higher for cues that predicted large rewards than for cues predicting small rewards (Figure S1B) indicating that mice had learnt the CS–US associations.

**Figure 1.**
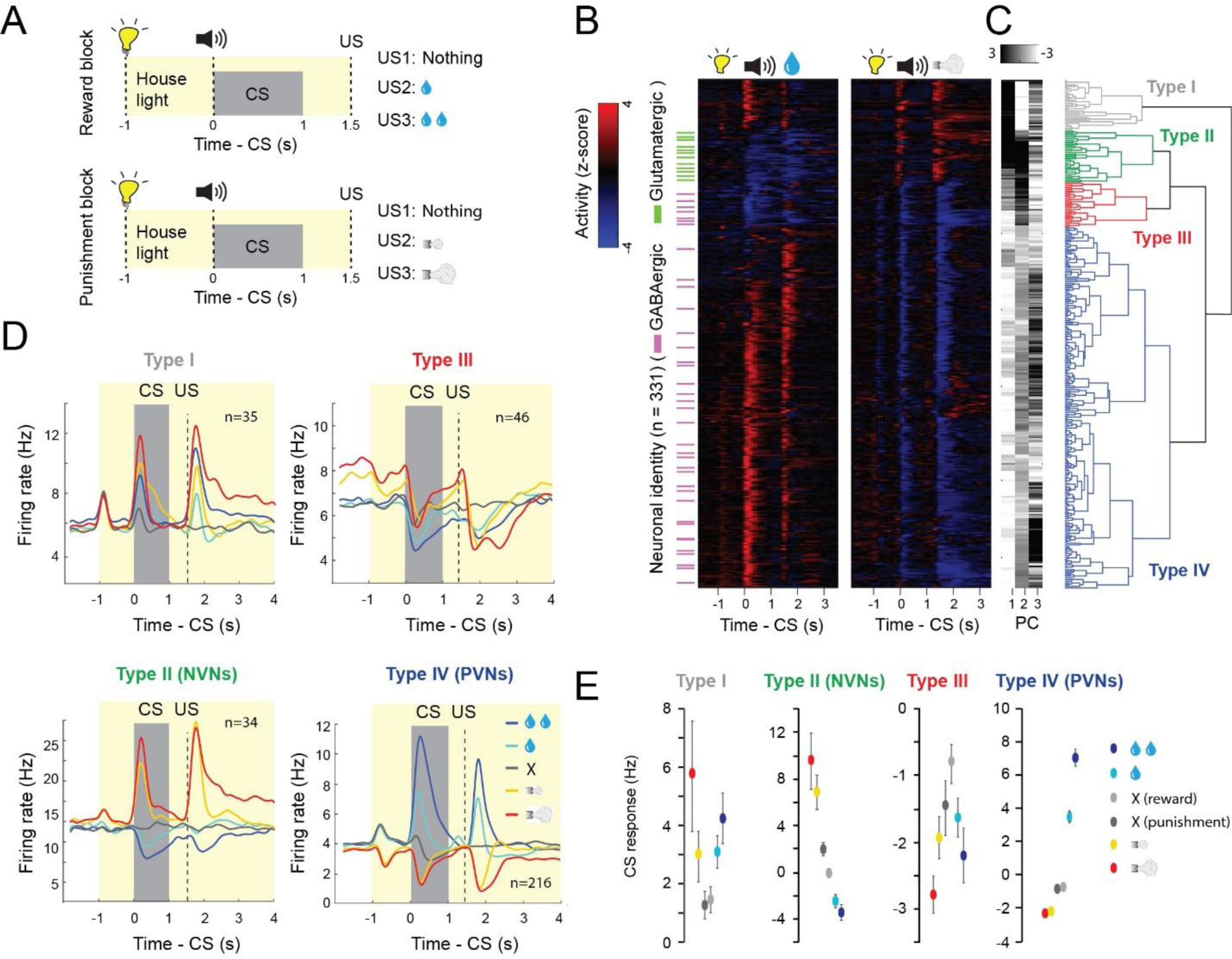
Separate VP populations opposingly encode motivational value and salience. **(A)** Illustrations of experimental design of the reward and punishment classical conditioning tasks. **(B)** Z-score activity plots of the responses of all neurons recorded in the reward and punishment tasks. Red, increase from baseline; blue, decrease from baseline; each row represents one neuron. Green and purple dashes indicate neurons that were optogenetically tagged as being glutamatergic and GABAergic, respectively. **(C)** First three principle components (PC) and hierarchical clustering dendrogram showing the relationship of each neuron within the four clusters. **(D)** Average firing rates of the four types of neurons in the reward and punishment blocks, shown as spike density functions (n=331 from 6 mice). **(E)** Average CS response magnitude in the reward and punishment blocks for each of the four types of neurons. All comparisons between the average CS responses were significant (at least *P*<0.05; Wilcoxon signed-rank test) except between the two neutral cues in Type I, III and IV neurons. There was also no significant difference between the CS response predicting large and small punishments in Type III neurons. Data in D are presented as mean ± s.e.m.

We recorded the activity of VP neurons (n = 331 neurons / 6 mice; 55 ± 15 per mouse) in *Vglut2-Cre;Ai40D* (n = 2) or *Gad2-Cre;Ai40D* (n = 4) mice, in which glutamatergic or GABAergic VP neurons could be optogenetically tagged due to the expression of the light-sensitive proton pump archaerhodopsin (ArchT) (see Methods) (Figure S2A-S2D). Hierarchical clustering revealed that there were four functional classes of VP neurons (Figure 1B, 1C). All identified glutamatergic neurons belonged to one functional cluster (type II). These neurons were activated by punishment-predictive CSs and punishments, and inhibited by reward-predictive CSs and rewards; we will refer to these as negative valence neurons (NVNs) (Figure 1B-1D). Two other clusters (type I, IV) contained identified GABAergic neurons (Figure 1B, 1C). Type IV neurons were activated by reward-predictive CSs and rewards, and inhibited by punishment-predictive CSs and punishments; we’ll refer to these as positive valence neurons (PVNs) (Figure 1B-1D). For both the negative (type II) and positive (type IV) value neurons, the magnitude of the CS responses was graded, reflecting the expected magnitude of reward or punishment (Figure 1E). We conclude that these neurons bi-directionally and oppositely encode the positive or negative valences of expected and actual outcomes.

In contrast to these two valence encoding populations, the two other populations appear to encode stimulus saliency as they were excited (type I) or inhibited (type III) by both rewards and punishments or the cues that predict these USs (Figure 1B-1E). Type I neurons were never identified as either glutamatergic or GABAergic and resemble cholinergic basal forebrain neurons that have been described (Hangya et al., 2015). Type III neurons that were inhibited by salient stimuli were identified as GABAergic (Figure 1B-1D).

Over the course of learning, the responses of both PVNs and NVNs to CS increased while their response to positive and negative US decreased, respectively (Figure 2A; Figure S3A-S3E). This reduction in US response in VP neurons is reminiscent of the temporal backpropagation seen in reward prediction error coding dopamine neurons (Cohen et al., 2012; Pan et al., 2005). To examine if these two populations encode reward and punishment prediction errors, we analysed their responses to the neutral cue following the house light. In our task the house light in each block predicts the possibility for reward or punishment, so in each case the delivery of the neutral cue represents an outcome that is worse or better than expected. NVNs were not significantly modulated when a neutral cue was presented in a punishment block, nor were they modulated when the neutral cue was presented in the reward block (Figure S3F-S3G), indicating that these neurons do not respond when an outcome is better or worse than expected. In contrast, PVNs displayed a decrease in firing relative to baseline when the neutral cue was presented in a reward block (Figure 2B-2C; Figure S3H-S3I). This indicates that they responded when an outcome was worse than expected. In the punishment block PVNs did not display an increase in firing when the neutral cue was delivered (Figure S3H-S3I). This suggests that PVNs selectively respond to reward but not punishment prediction errors. To confirm that this decrease in firing was associated with a worse than expected outcome, we omitted an expected reward in 10% of trials. Again, PVNs displayed a decrease in firing relative to baseline (Figure 2D-2E). These properties of the PVNs are similar to those of the reward prediction error-coding dopamine neurons (Cohen et al., 2012; Pan et al., 2005; Schultz et al., 1997).

**Figure 2.**
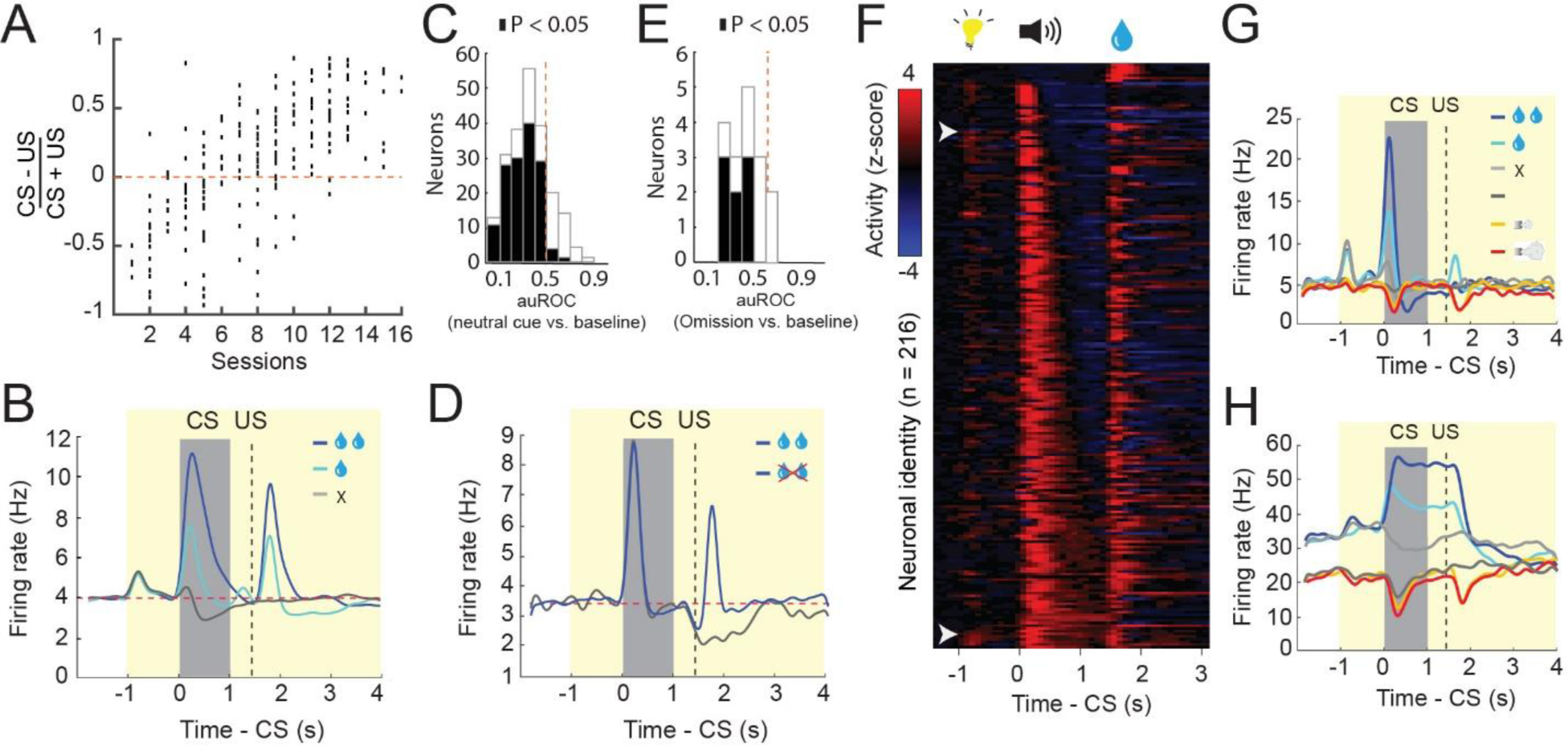
VP neuron responses are modulated by expectation. **(A)** CS-US (reward) response index for all type IV neurons across different stages of training (*r*^2^ = 0.69, P < 0.001 by linear regression). **(B)** Average firing rates of type IV neurons in the reward block, shown as spike density functions. **(C)** auROC analysis of difference in firing rate between baseline and neutral cue presentation (n = 221 from 6 mice). Filled bars, *P*<0.05, *t*-test. **(D)** Average firing rates of type IV neurons in response to reward omission, shown as spike density functions. **(E)** auROC analysis of difference in firing rate between baseline and reward omission trials (n = 15 from 2 mice). Filled bars, *P*<0.05, *t*-test. **(F)** Z-score activity plots of the responses of all type IV neurons sorted for the duration of their CS response. **(G, H)** Two example neurons showing phasic (G) and sustained (H) responses to reward predicting cues.

While prediction error-coding dopamine neurons respond phasically to reward predicting cues, the duration of the CS response in VP PVNs was variable (Figure 2F-2H). Indeed, a number of PVNs had a sustained CS response that lasted till the onset of the US (Figure 2F-2H). These sustained responsive neurons are similar to the VP neurons that have been reported to encode state values (Tachibana and Hikosaka, 2012). However, when sorted for the duration of the CS responses there did not appear to be two populations of PVNs (phasic and sustained) (Figure 2F), nor did our hierarchical clustering identify two sub-clusters within the population; rather the population appeared to represent a continuum as has been previously reported (Richard et al., 2016; Richard et al., 2018).

As the VP has been intricately linked to motivation, we examined if the value coding depended on the motivational state of the mice. In the reward sessions of the Pavlovian task, mice showed vigorous licking response starting from CS onset and lasting until well beyond the delivery of water in the early trials, a period when mice were thirsty, but dramatically reduced licking towards the end of a session (Figure 3A-3B). This decrease in licking presumably reflects a reduction in the motivation to pursue water as the mice were sated. Remarkably, the excitatory CS responses of the PVNs, which were prominent in ‘thirsty trials’, completely disappeared and inverted in ‘sated trials’ (Figure 3A-3D). By contrast, the NVNs were differently modulated by animals’ thirsty state. While in thirsty trials these neurons were inhibited by the CS predicting water delivery, in stated trials they were excited by the same CS (Figure 3A-3D). Notably, thirst also strongly modulated the baseline activities of PVNs and NVNs, such that both populations showed markedly lower baseline activities in sated trials than in thirsty trials (Figure 3C-3D).

**Figure 3.**
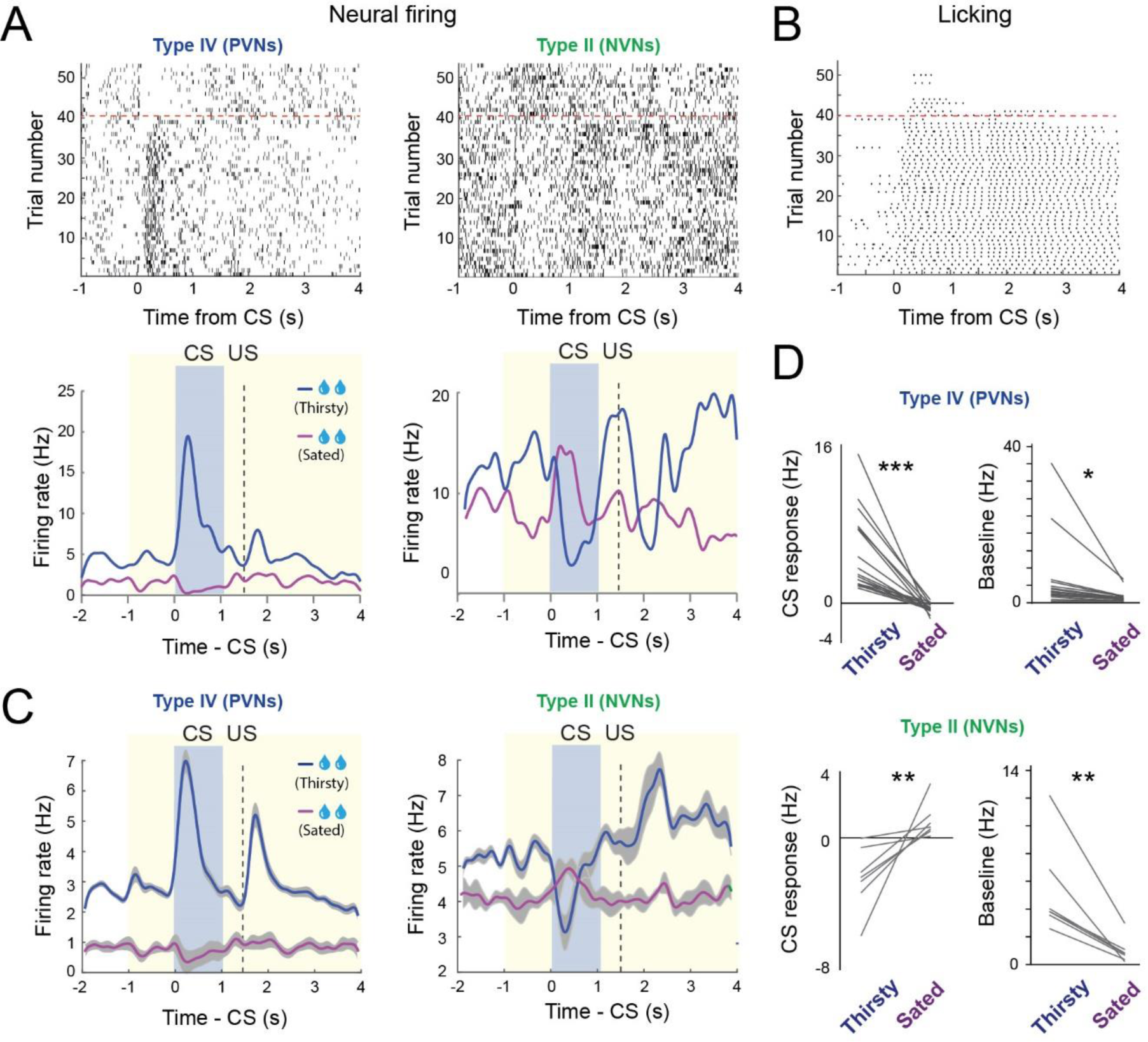
The response of VP neurons to reward predicting CS depends on the internal motivational state. **(A)** Top: raster plots showing the neural activity of a PVN (left) and a NVN (right) during large reward trials. Bottom: spike density plots showing the average response of the corresponding two neurons when the mouse was thirsty or sated. **(B)** Raster plot showing the licking behavior in the same behavioral session. **(C)** Average spike density plots showing the activity of PVNs (n = 19, from 2 mice) and NVNs (n = 7, from 2 mice) in thirsty and sated trials. **(D)** Left: graphs showing the average CS response of PVNs (top) and NVNs (bottom) when mice were thirsty or sated. Right: graphs showing the baseline firing rates of PVNs (top) and NVNs (bottom) when mice were thirsty or sated (*** *P* < 0.001, ** *P* < 0.01, * *P* < 0.05; paired *t*-test). Data in C are presented as mean ± s.e.m. (shaded areas).

Thus, both changes in the predicted value of the environment and the animal’s internal state differentially modulates the activities in PNVs and NVNs, as reflected in both the transient cue-induced responses and the tonic baseline activities in these neurons. These results point to the possibility that PNVs and NVNs differentially and potentially opposingly contribute to the generation of incentive and aversive motivation.

### PVNs and NVNs opposingly and cooperatively regulate motivation

To investigate how PVNs and NVNs might influence the motivation to approach or avoid, we virally expressed channelrhodopsin (ChR2) or ArchT in GABArgic or glutamatergic VP neurons. We found that optogenetic activation of GABAergic or glutamatergic VP neurons induced real-time place preference (RTPP) or aversion (RTPA), respectively (Figure S4A-S4B). Furthermore, optogenetic activation of GABAergic VP neurons also supported self-stimulation (Figure S4C). In contrast, optogenetic inhibition of these neurons induced neither RTPP nor RTPA (Figure S4D-S4E). These results, which are largely consistent with previous findings (Faget et al., 2018; Knowland et al., 2017a; Tooley et al., 2018a), suggest that activation of PVNs or NVNs are innately appetitive or aversive, respectively.

To understand how the PVNs and NVNs contribute to motivated behaviour, we first designed a reward-and-punishment conflict task (or ‘conflict task’), in which incentive value can be modulated by either changing reward size or by introducing punishment (Figure 4A). Specifically, in the reward block of this task, mice needed to lick during a choice window following a CS in order to obtain a water reward, whereas in the conflict block, licking during the choice window led to simultaneous delivery of a water reward and an air-puff in each trial (Figure 4A). In both blocks, different CSs predicted rewards of different sizes. We found that the licking probability during the choice window (i.e., choice probability) increased as reward size increased, as would be expected from the associated increase in incentive value. In contrast, choice probability decreased when the reward was paired with an air-puff, as would be expected from a decrease in incentive value (Figure 4B).

**Figure 4.**
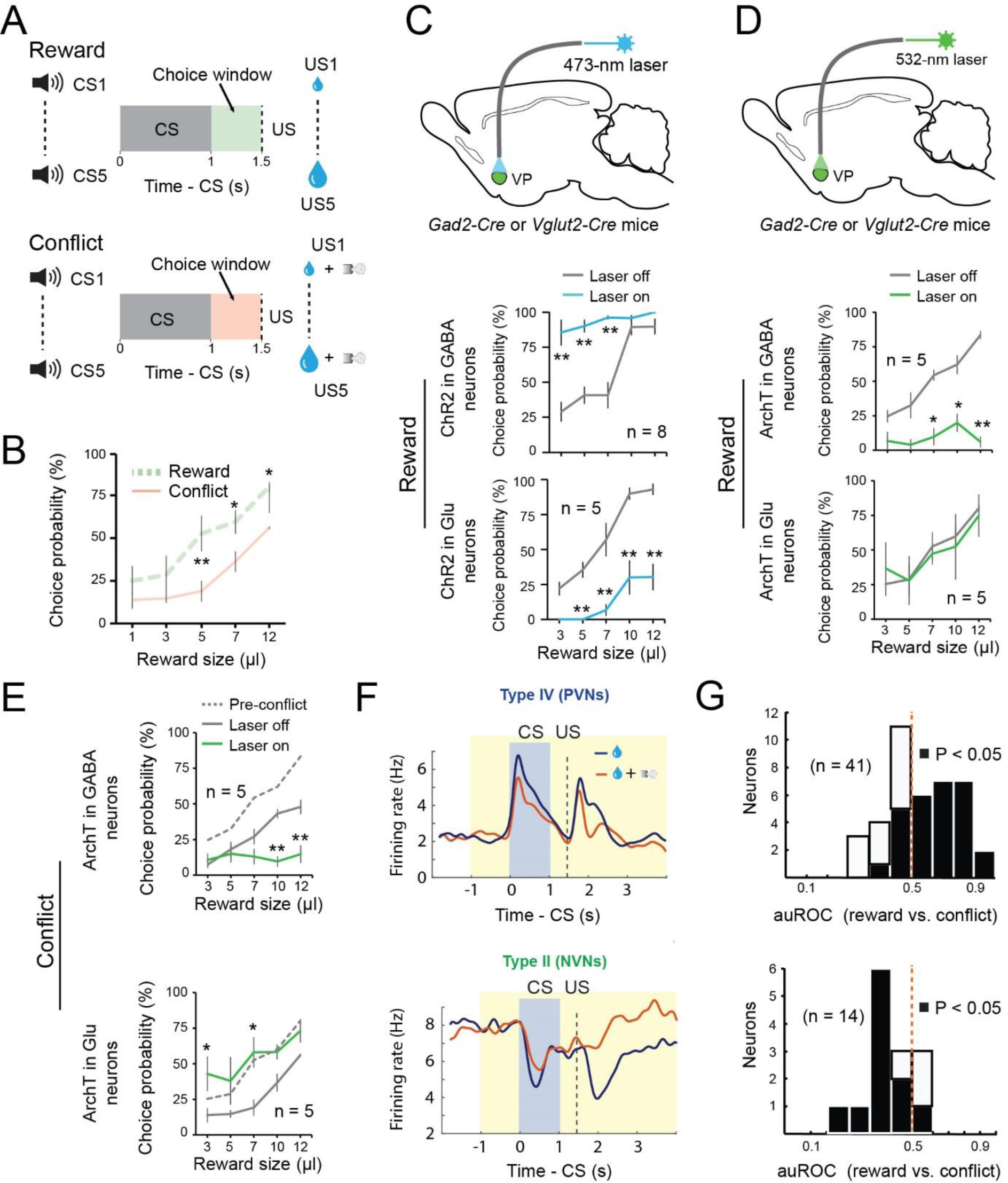
The balance of activity between GABAergic and glutamatergic VP neurons controls reward seeking during motivational conflict. **(A)** Schematics of the experimental design. **(B)** Motivational conflict reduced reward seeking (F_(1, 9)_ = 35.19, p < 0.0001). **p < 0.01, *p < 0.05, two-way ANOVA followed by Tukey’s test. **(C)** Top: a schematic of the approach. Middle: activation of GABAergic VP neurons increased reward seeking (F_(1,15)_= 92.32, p < 0.001). Bottom: activation of glutamatergic VP neurons decreased reward seeking (F_(1, 7)_ = 108.68, p < 0.001). **P < 0.01, two-way ANOVA followed by Tukey’s test. **(D)** Top: a schematic of the approach. Middle: inhibition of GABAergic VP neurons decreased reward seeking (F_(1, 9)_ = 50.37, p < 0.0001). Bottom: inhibition of glutamatergic VP neurons had no effect on reward seeking (F_(1, 9)_ = 0.055, p = 0.82). **P < 0.01, *P < 0.05, two-way ANOVA followed by Tukey’s test. **(E)** Top: inhibition of GABAergic VP neurons further decreased reward seeking in the conflict task (F_(1, 9)_ = 27.60, p < 0.0001). Bottom: inhibition of glutamatergic VP neurons increased reward seeking to pre-conflict levels (F_(1, 9)_ = 50.37, p < 0.0001). **p < 0.01, *p <0.05, two-way ANOVA followed by Tukey’s test. **(F)** Average response of PVNs (top; n = 41, from two mice) or NVNs (bottom; n = 14, from two mice) in reward or conflict trials, shown as spike density plots. **(G)** auROC analysis of difference in the CS response during reward and conflict trials. Top: PVNs (n = 41 from two mice); bottom: NVNs (n = 14, from two mice). Filled bars, *P*<0.05, *t*-test. Data are presented as mean ± s.e.m.

To test how the different classes of VP neurons influence choice probability, we optogenetically activated or inhibited these neurons, as described above, during the time window between CS onset and US onset (which covered the choice window) in 20% of randomly selected trials in the conflict task. Notably, activation of GABAergic VP neurons increased the probability that mice would lick in the choice window, in particular when the CS predicted a small reward (i.e., when the motivation to lick was low) (Figure 4C); inhibition of these neurons decreased the probability that mice would lick on a given trial, even if the CS predicted a large reward (i.e., when the incentive to lick was high) (Figure 4D). By contrast, activation of glutamatergic VP neurons decreased the probability that mice would lick on a given trial (Figure 4A). Although inhibition of the glutamatergic VP neurons had no effect on the behavior in reward blocks (Figure 4D), inhibition of these neurons in the conflict blocks increased the probability that mice would lick in a trial (Figure 4E). In control experiments we verified that light illumination alone in the VP had no effect on animal behavior in the conflict task (Figure S5A-S5B). These results suggest that GABAergic VP neurons play an essential role in generating or regulating incentive value. By contrast, glutamatergic VP neurons are not needed for reward seeking but are needed to constrain reward seeking when there is cost (punishment) associated with the action.

To determine how the endogenous activities of these VP neurons might be modulated by the ‘conflict’, we recorded VP neurons in mice performing a modified version of the conflict task (see Methods). We found that, in the conflict trials as compared to the reward-only trials, the excitatory CS responses of PVNs were reduced, whereas the activity of the NVNs was higher in the conflict trials (Figure 4F-4G). These results indicate that perceived risk associated with licking in the conflict task suppresses and increases (or disinhibits), respectively, the responses of PVNs and NVNs to reward cues. These results, together with those from optogenetic manipulations, suggest that the balance between PVN’s and NVN’s activity sets the motivation to seek the reward when there is a motivational conflict.

While the NVNs do not play a direct role in driving reward-seeking behavior, we hypothesized that they may drive behavior in an aversive context. To determine how PVNs and NVNs contribute to behaviour in an appetitive or aversive context, we designed a “run-for-water” (RFW) task and a “run-to-avoid-air-puff” (RTAA) task, respectively. In the RFW task, mice needed to run in response to a CS in order to obtain a water reward, whereas in the RTAA task mice had to run in response to a CS in order to avoid an air-puff (Figure S6A-S6D). Once mice learned the tasks, we optogenetically activated or inhibited GABAergic or glutamatergic neurons in the VP, as described above, during the time window between CS onset and US onset in 20% of randomly selected trials (Figure 5A-5F; Figure S7). We found that, in the RFW task, activating and inhibiting GABAergic neurons promoted and abolished running, respectively (Figure 5B, 5C, 5E; Figure S8A-S8B). In contrast, activating glutamatergic neurons decreased the velocity in the RFW task, while inhibiting these neurons did not affect running (Figure 5B, 5C, 5E; Figure S8A-S8B). In stark contrast to the RFW task, in the RTAA task, activating GABAergic neurons completely abolished running, whereas inhibiting these neurons had no effect (Figure 5B, 5D, 5F; Figure S8A-S8B); moreover, activating and inhibiting glutamatergic neurons promoted and abolished running, respectively (Figure 5B, 5D, 5F; Figure S8A-S8B). Thus, while GABAergic neurons promote running in a reward-seeking task, activating them suppresses running in a punishment-avoidance task. In the opposite manner, glutamatergic neurons are essential for promoting running to avoid punishment, but suppress the same behavior when it is being performed to obtain a reward. These results suggest that the motivational context switches the role that each population plays in behavior; in a reward-seeking context PVNs drive the behaviour while NVNs constrain the behaviour. In an aversive context these roles reverse, with the NVNs driving avoidance behavior and the PVNs actively constraining the avoidance.

**Figure 5.**
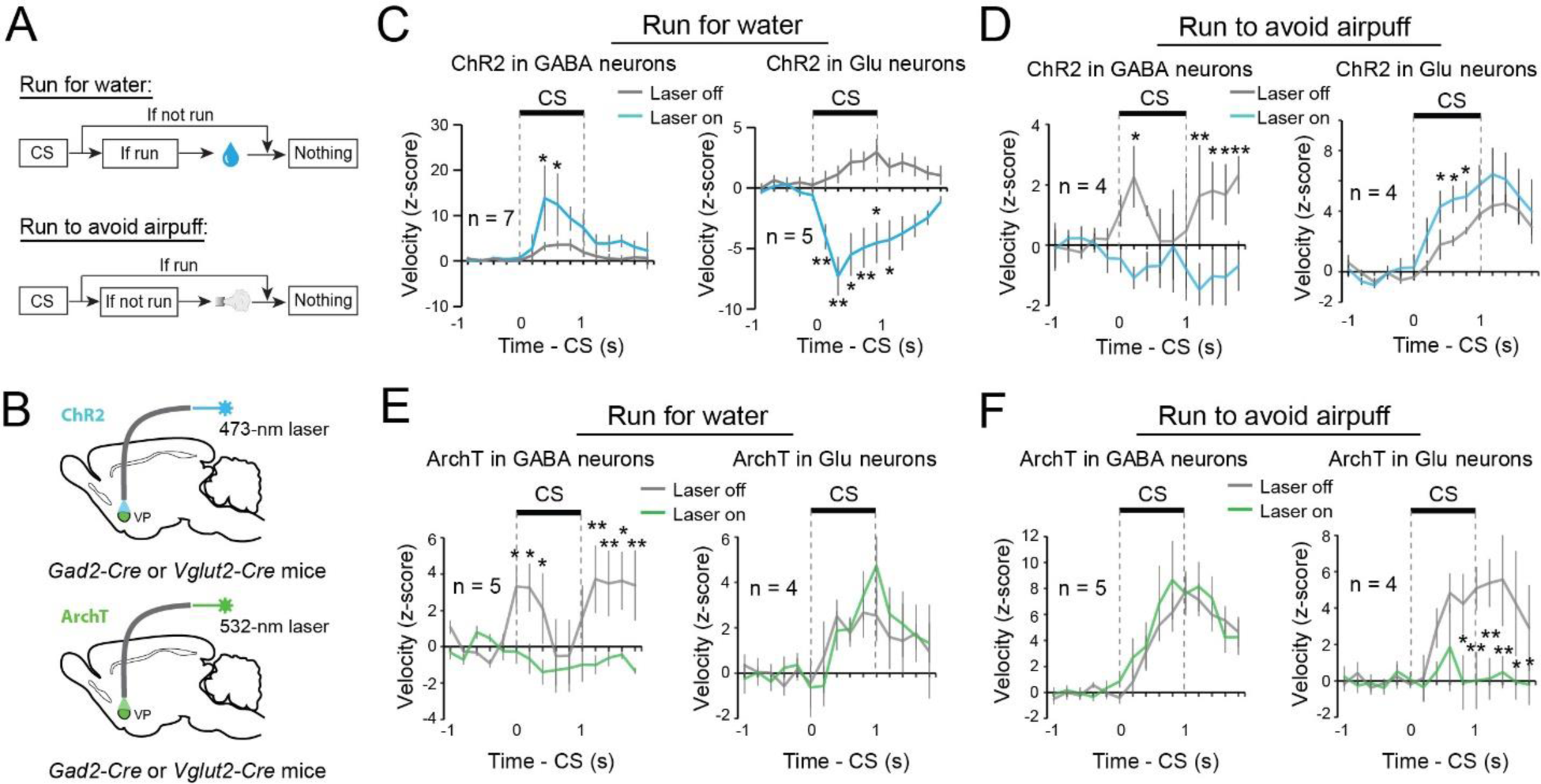
GABAergic and glutamatergic VP neurons switch roles in controlling actions when motivational context changes. **(A)** Schematics of the experimental design. **(B)** Schematics of the approach. **(C)** Left: activation of GABAergic VP neurons increased running for water reward (F_(1, 13)_ = 7.90, p = 0.0055). Right: activation of glutamatergic VP neurons decreased running for water reward (F_(1, 9)_ = 132.73, p <0.001). **P < 0.01, *P < 0.05, two-way ANOVA followed by Tukey’s test. **(D)** Left: activation of GABAergic VP neurons decreased running to avoid air puff (F_(1,7)_ = 18.76, P < 0.0001). Right: activation of glutamatergic VP neurons increased running to avoid air puff (F_(1,7)_ = 11.61, P = 0.0010). **P < 0.01, *P < 0.05, two-way ANOVA followed by Tukey’s test. **(E)** Left: inhibition of GABAergic VP neurons decreased running for water reward (F_(1,9)_ = 29.283, p < 0.0001). Right: inhibition of glutamatergic VP neurons had no effect on running for water reward (F_(1, 7)_ = 0.30, p = 0.59). **P < 0.01, *P < 0.05, two-way ANOVA followed by Tukey’s test. **(F)** Left: inhibition of GABAergic VP neurons had no effect on running to avoid air puff (F_(1,9)_ = 1.30, p = 0.26). Right: inhibition of glutamatergic VP neurons decreased running to avoid air puff (F_(1, 7)_ = 22.06, p <0.001). **P < 0.01, *P < 0.05, two-way ANOVA followed by Tukey’s test. Data are presented as mean ± s.e.m.

### PVNs and NVNs act via the VP–LHb pathway

In order to explore how downstream structures integrate the activity of PVNs and NVNs, we first looked at the projections of GABAergic and glutamatergic VP neurons. As has previously been reported, both GABAergic and glutamatergic neurons receive input from areas such as the nucleus accumbens, prefrontal cortex and basolateral amygdala (Figure S9A-S9G) and project to qualitatively the same structures, including the ventral tegmental area, lateral hypothalamus, lateral habenula (LHb) and rostromedial tegmental nucleus (Root et al., 2015; Tooley et al., 2018b). Each population of VP neurons may synapse onto different cell types within these areas or individual neurons may receive opposing inputs from both VP populations. To explore this, we choose one projection target, the LHb. The VP projects to the medial portion of the LHb (Figure S10) (Herkenham and Nauta, 1977; Root et al., 2015), which in turn projects to the dorsal raphe (DR) (Herkenham and Nauta, 1979; Quina et al., 2015). The DR also has neurons that encode the motivational value (Cohen et al., 2015) and the behavioural effects of serotonin, like those reported here, also depend on the state of the environment (Seo et al., 2019). Retrograde tracing combined with single molecule in situ hybridization confirmed that both GABAergic and glutamatergic VP neurons project to the LHb (Figure S11). In addition, using optogenetics combined with electrophysiology in acute slices, we recorded from LHb neurons that were retrogradely labeled from the DR. Every neuron that was recorded received GABAergic or glutamatergic VP input, suggesting that individual DR-projecting LHb neurons receive inputs from both VP populations (Figure S12A-S12D).

To determine the behavioural effect of the VP→LHb pathway, we optogenetically activated GABAergic or glutamatergic VP axon terminals in the LHb. Activation of the GABAergic (Gad2^VP→LHb^) or glutamatergic (Vglut2^VP→LHb^) inputs to the LHb induced RTPP or RTPA, respectively (Figure S13A, S13B; Figure S10). These results confirmed that both the GABAergic and the glutamatergic VP neurons send substantial projections to the LHb that can differentially influence animal behavior, consistent with previous findings (Faget et al., 2018; Knowland et al., 2017a; Tooley et al., 2018a).

Next, we assessed the roles of Gad2^VP→LHb^ and Vglut2^VP→LHb^ in the behaviors driven by incentive or aversive value, as described above. In the reward and conflict tasks, we found that optogenetically inhibiting Gad2^VP→LHb^ decreased the choice probability in the reward-only block (Figure 6A, 6B). This manipulation did not reduce choice in the conflict block, likely due to a “floor effect” as this group of mice were highly sensitive to the punishment (Figure 6C). By contrast, inhibiting Vglut2^VP→LHb^ did not affect choice in the reward-only block, but increased the choice probability in the conflict block (Figure 6B, 6C).

**Figure 6.**
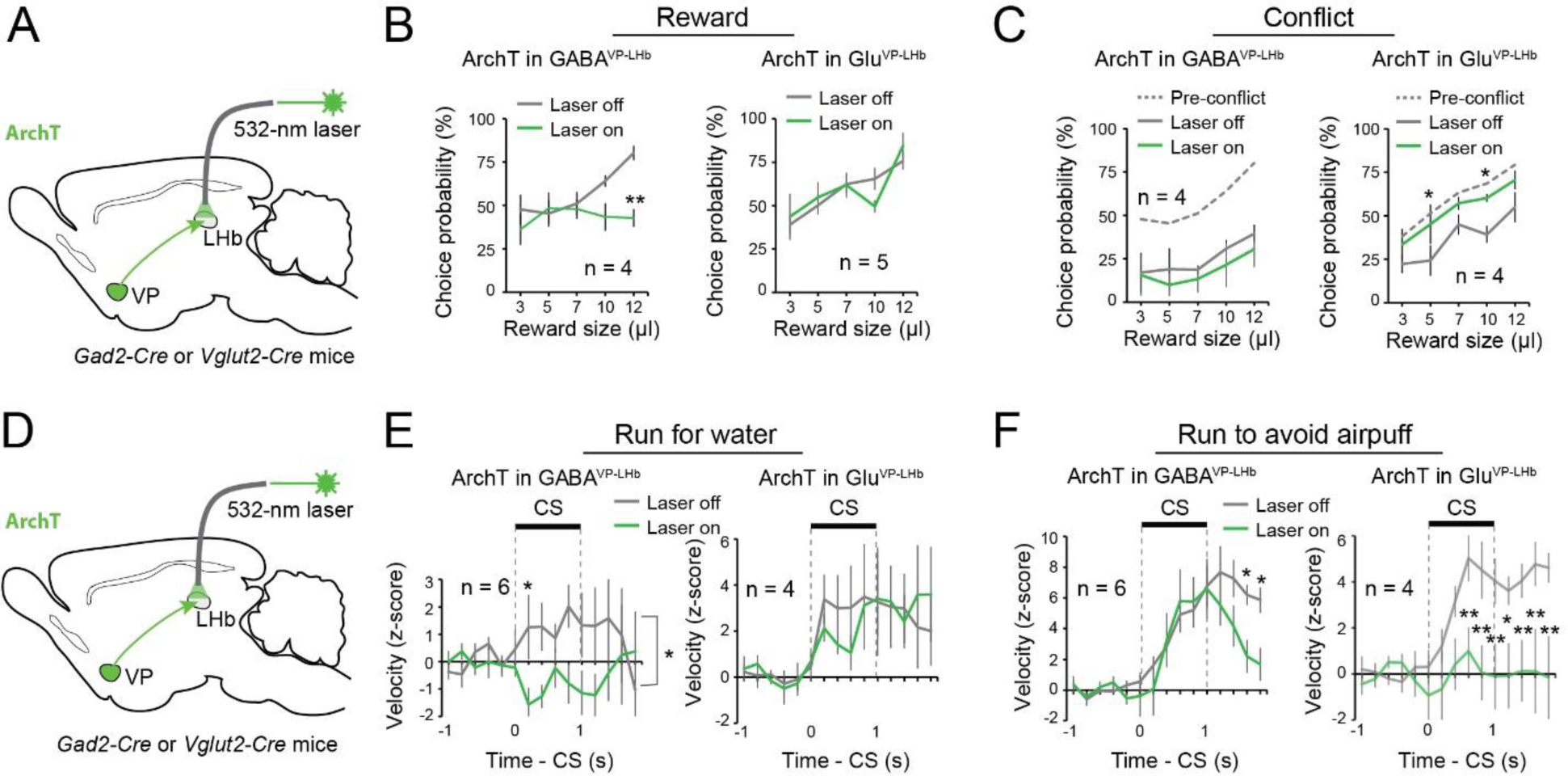
GABAergic and glutamatergic VP neurons act via the VP-LHb pathway. **(A)** A schematic of the approach. **(B)** Left: inhibition of GABAergic^VP→LHb^ decreased reward seeking (F(_1, 7)_ = 9.55, p = 0.0043). Right: inhibition of glutamatergic^VP→LHb^ had no effect on reward seeking (F_(1, 9)_ = 0.0041, p = 0.95). **P < 0.01, two-way ANOVA followed by Tukey’s test. **(C)** Left: inhibition of GABAergic^VP→LHb^ did not further decrease reward seeking in these mice in the conflict task (F_(1, 7)_ = 1.39, p = 0.25). Right: inhibition of glutamatergic^VP→LHb^ increased reward seeking to pre-conflict levels (F_(1, 7)_ = 13.17, p = 0.0010). **P < 0.01, *P < 0.05, two-way ANOVA followed by Tukey’s test. **(D)** A schematic of the approach. **(E)** Left: inhibition of GABAergic^VP→LHb^ decreased running for water reward (F_(1, 11)_ = 8.56, p = 0.004). Right: inhibition of glutamatergic^VP→LHb^ had no effect on running for water reward (F_(1, 7)_ = 0.060, p = 0.81). *P < 0.05, two-way ANOVA followed by Tukey’s test. **(F)** Left: inhibition of GABAergic^VP→LHb^ had no effect on running to avoid air puff during the cue, although it induced an earlier termination of the running response after the cue (F_(1, 11)_ = 5.57, p = 0.020). Right: inhibition of glutamatergic^VP→LHb^ decreased running to avoid air puff (F_(1, 9)_ = 40.72, p < 0.0001). **P < 0.01, *P < 0.05, two-way ANOVA followed by Tukey’s test. Data are presented as mean ± s.e.m.

In the RFW and RTAA tasks (see Figure 5), inhibiting Gad2^VP→LHb^ abolished running for water (Figure 6D, 6E), but only slightly reduced running to avoid air-puff (Figure 6F). By contrast, inhibiting Vglut2^VP→LHb^ did not affect running for water (Figure 6E), but abolished running to avoid air-puff (Figure 6F). Of note, inhibition of either pathway did not induce any obvious effects in the RTPP or RTPA task (Figure S13C, S13D). Furthermore, control experiments demonstrated that the behavioral effects we observed were not induced by light illumination per se (Figure S14A-S14D). Together, these results are consistent with the optogenetic manipulations of cells in the VP (Figure 4 & 5), and suggest that the distinct behavioral roles of the GABAergic and glutamatergic VP neurons are, at least in part, mediated by their projections to the LHb.

## DISCUSSION

The decision to approach or avoid depends on the situation in the environment and the internal state of the animal. Here we show that separate populations of VP neurons drive behaviour in different motivational contexts and their activity is differentially regulated by both the valence of the environment and the internal state of the animal. GABAergic neurons are necessary for driving reward-seeking behaviour in positive motivational context, while glutamatergic neurons are needed for driving avoidance behaviour in an aversive motivational context. Previous work had shown that the VP is important for reward-seeking and punishment avoidance (Root et al., 2015; Smith et al., 2009; Stephenson-Jones, 2019; Wulff et al., 2018). Our results now show that these different functions are driven by separate populations of VP neurons.

In the natural world, reward seeking is associated with certain costs, such as the effort needed to obtain a reward, the risk of punishment or the presence of threats (Ydenberg, 1986). Our results show that glutamatergic neurons are needed to constrain reward seeking when there is a risk associated with seeking reward. In line with this a recent study showed that glutamatergic VP neurons are needed to constrain reward seeking when effort is required to obtain a reward, or to limit reward seeking when the reward has been devalued (Tooley et al., 2018b). In light of this finding our results suggest that glutamatergic neurons represent the “costs” associated with seeking-reward and work to balance the incentive value represented by the GABAergic neurons. In the opposing manner the internal state of an animal, such as thirst may encourage animals to take a risk and reduce the likelihood an animal would avoid a threating environment. Our results now suggest that GABAergic neurons can constrain avoidance. Together our results suggest that the balance between GABAergic and glutamatergic VP activity sets the motivation for approach and avoidance.

If the balance between these populations sets the behavioural response in different motivational contexts, how then do downstream neurons read out this balance? Our results suggest one mechanisms by which this is achieved, i.e., individual neurons downstream of the VP receive input from both GABAergic and glutamatergic VP neurons. In this way individual neurons can integrate the excitatory and inhibitory input from the VP, with the balance between these drives determining the activity of the postsynaptic neuron. While other patterns of integration likely exist, individual neurons in the VTA also appear to integrate the excitatory and inhibitory input from the VP, as more than half of VTA neurons receive direct GABAergic or glutamatergic VP input (Faget et al., 2018). This suggests that as predicted from their common projection pattern (Faget et al., 2018), the different VP populations have an opposing influence on common downstream targets.

If downstream neurons integrate input from the different VP populations, they should be activated or inhibited in opposing motivational contexts. In line with this a recent study has shown that GABAergic and serotoninergic neurons in the dorsal raphe are selectively activated and drive movement in high threat environments, but are inhibited and suppress movement in a rewarding environment (Seo et al., 2019). Interestingly, the VP innervates the medial portion of the LHb which projects to the dorsal raphe (Herkenham and Nauta, 1977, 1979). As with neurons in the VP the firing rate of the serotonergic neurons is modulated on a short and long timescale by the predicted value and utility of a stimulus (Cohen et al., 2015). This suggests the contextual dependence of raphe activation may be driven by a switch in the balance between GABAergic and glutamatergic VP activity.

Previous *in vivo* recording data has shown that VP neurons encode a number of variables related to motivation such as the expected reward value (Tachibana and Hikosaka, 2012; Tian et al., 2016), state and action values (Ito and Doya, 2009; Saga et al., 2017; Tachibana and Hikosaka, 2012) as well as reward prediction error (Tian and Uchida, 2015). In line with these findings the activity of a subset of our value coding GABAergic neurons could be characterized as encoding reward prediction error or state value. Despite this, our data suggest that value coding GABAergic neurons represent one continuous variable with different neurons responding to motivationally relevant cues with different durations. An alternative variable has been suggested to account for the activity pattern of VP neurons, this is incentive value, or the degree to which stimuli have the ability to activate motivational states (Richard et al., 2016; Richard et al., 2018; Smith et al., 2009). This variable may better account for the GABAergic VP activity that we describe here, as it relates both the external stimulus value as well as the internal state of the animal (Berridge, 2012). This is important as previous recordings, as does our study, show that the activity of VP neurons depends on the internal motivational state of the animal as well as the external environment (Smith et al., 2009; Tindell et al., 2006). In light of these findings we propose that value coding GABAergic and glutamatergic neurons encode incentive and aversive value, respectively.

The decision to approach or avoid depends on the situation in the environment and the internal state of the animal. Here we show that two populations of VP neurons are critical for driving approach and avoidance behaviour. Each of these populations is differentially regulated by both the internal state and the predicted motivational value. These results indicate that the VP is a critical area where information about the internal state and the environmental context is combined to determine the overall behavioural strategy to either approach or avoid.

## ACKNOWLEDGEMENTS

We thank Dr. Thomas Mrsic-Flogel for critically reading the manuscript, Dr. Z. Josh Huang for providing mouse strains, Ga-Ram Hwang and Dylan Rebolini for technical assistance, and members of the Li laboratory for helpful discussions. This work was supported by grants from NARSAD (M.S., S.A., B.L.), the National Institutes of Health (NIH) (F32MH113316, C.B.R; R01MH101214, R01MH108924, R01NS104944, B.L.), Human Frontier Science Program (RGP0015/2016, B.L.), the Stanley Family Foundation (B.L.), Simons Foundation (344904, B.L.), Wodecroft Foundation (B.L.), the Cold Spring Harbor Laboratory and Northwell Health Affiliation (B.L.) and Feil Family Neuroscience Endowment (B.L.).

## AUTHOR CONTRIBUTIONS

M.S., C.B.R and B.L. designed the study. M.S. and C.B.R conducted the majority of the experiments and analyzed data. S.A. performed the patch clamp recording experiments. A.F. performed the RNAscope experiments. C.F.H. performed the rabies tracing experiments. M.S. and B.L. wrote the paper with inputs from all authors.

## METHODS

### Animals

All procedures were approved by the Institutional Animal Care and Use Committee of Cold Spring Harbor Laboratory (CSHL) and conducted in accordance to the United States’ National Institutes of Health guidelines. Mice were housed under a 12 h light-dark cycle (8 a.m. to 8 p.m. light). All behavioural experiments were performed during the light cycle. All mice had free access to food, but water was restricted for mice used in certain behavioural experiments. Free water was provided on days with no experimental sessions. Male and female mice 2-4 months of age were used in all experiments. No differences were observed in the behavior of male or female mice during the behavioural tasks or with the optogenetic manipulations (see below). All animals were randomly allocated to the different experimental conditions used in this study. The *Vglut2-Cre* (*Slc17a6^tm2(cre)Lowl^/J*, stock #016963 from Jackson Laboratory, Bar Harbor, Maine, USA), *GAD2-IRES-Cre* (from Dr. Z. Josh Huang, CSHL, available from Jackson Laboratory, *Gad2^tm2(cre)Zjh^/J*, stock # 010802), *Ai40D* (*Gt(ROSA)_26Sor_^tm40.1(CAG-aop3/GFP)Hze/J^*), stock #021188 from Jackson Laboratory), *Rosa26-stop^flox^-tTA* (stock #012266 from Jackson Laboratory) (Li et al., 2010; Penzo et al., 2015) mouse strains have all been previously characterized. All mice were bred onto a C57BL/6J background.

### Viral vectors

All adeno-associated viruses (AAV) were produced by the University of North Carolina vector core facility (Chapel Hill, North Carolina, USA) or the University of Pennsylvania vector core (Pennsylvania, USA) and have previously been described: AAV9-Ef1a-DIO-hChR2(H134R)-eYFP, AAV9-CAG-FLEX-ArchT-GFP, AAV9-Ef1a-DIO-eYFP and AAV-TRE-hGFP-TVA-G. The EnvA-pseudotyped, protein-G-deleted rabies-EnvA-SAD-ΔG-mCherry virus (Miyamichi et al., 2011) was produced by the Viral Vector Core Facility at Salk Institute. All viral vectors were aliquoted and stored at −80 °C until use.

### Stereotaxic surgery

Mice were anesthetized with isoflurane inhalant gas (3%) in an induction chamber and positioned in a stereotaxic frame (myNeuroLab, Leica Microsystems Inc., Buffalo Grove, Illinois, USA). Isoflurane (1.5%) was be delivered through a facemask for anesthesia maintenance. Lidocaine (20 µl) was injected subcutaneously into the head and neck area as a local anaesthetic. For *in vivo* recordings, mice were implanted with a head-bar and a microdrive containing the recording electrodes and an optical fibre. Viral injections were performed using previously described procedures (Penzo et al., 2015) at the following stereotaxic coordinates: VP, 0.75 – 0.30 mm from bregma, 1.00 mm lateral from midline, and 5.10 mm ventral from cortical surface; LHb, –1.82 mm from bregma, 0.40 mm lateral from midline, and 2.46 mm ventral from cortical surface; and DR, −4.24 mm from bregma, 0 mm lateral from midline, and 3.00 mm ventral from cortical surface. During the surgical procedure, mice were kept on a heating pad and were brought back to their home-cage for post-surgery recovery and monitoring. Postoperative care included intraperitoneal injection with 0.3-0.5 ml of Lactated Ringer’s solution and Metacam (1-2 mg kg^−1^ meloxicam; Boehringer Ingelheim Vetmedica, Inc., St. Joseph, Missouri, USA) for analgesia and anti-inflammatory purposes. All AAVs were injected at a total volume of approximately 0.2 µl, and were allowed at least 4 weeks for maximal expression. For retrograde tracing of projection cells in the VP, CTB-555 (0.2 µl, 0.5% in phosphate-buffered saline (PBS); Invitrogen, Thermo Fisher Scientific, Waltham, Massachusetts, USA) was injected into the LHb or DR and allowed 5 days for sufficient retrograde transport.

### Immunohistochemistry

Immunohistochemistry experiments were performed following standard procedures. Briefly, mice were anesthetized with Euthasol (0.4 ml; Virbac, Fort Worth, Texas, USA) and transcardially perfused with 40 ml of PBS, followed by 40 ml of 4% paraformaldehyde in PBS. Coronal sections (50 μm) were cut using a freezing microtome (Leica SM 2010R, Leica). Sections were first washed in PBS (3 x 5 min), incubated in PBST (0.3% Triton X-100 in PBS) for 30 min at room temperature (RT) and then washed with PBS (3 x 5 min). Next, sections were blocked in 5% normal goat serum in PBST for 30 min at RT and then incubated with primary antibodies overnight at 4 °C. Sections were washed with PBS (5 x 15 min) and incubated with fluorescent secondary antibodies at RT for 1 h. After washing with PBS (5 x 15 min), sections were mounted onto slides with Fluoromount-G (eBioscience, San Diego, California, USA). Images were taken using a LSM 710 laser-scanning confocal microscope (Carl Zeiss, Oberkochen, Germany). The primary antibodies used were: chicken anti-GFP (1:1000, Aves Labs, catalogue number GFP1020, lot number GFP697986), rabbit anti-RFP (1:1000, Rockland, catalogue number 600-401-379, lot number 34135), and rabbit anti-Substance P (SP) (1:1000, ImmunoStar, catalog number 20064, lot number 1531001). Primary antibodies were incubated with appropriate fluorophore-conjugated secondary antibodies (1:1000, Life Technologies, Carlsbad, California, USA) depending on the desired fluorescence colour.

### Fluorescent *in situ* hybridization

Single molecule fluorescent *in situ* hybridization (ACDBio, RNAscope) was used to detect expression of *Gad2* and *Slc17a6* (*Vglut2*) in LHb-projecting VP neurons. Alexa-Fluor 555 Conjugate Cholera Toxin Subunit B (CTB555, Thermo Fisher Cat. No. C22843) was injected unilaterally into the LHb of wild-type adult mice. After 5 days, mice were decapitated and their brain tissue was first embedded in cryomolds (Sakura Finetek, Cat. No. 25608-924) filled with M-1 Embedding Matrix (Thermo Scientific, Cat. No. 1310) then quickly fresh-frozen on dry ice. The tissue was stored at −80°C until it was sectioned. 12 μm cryostat-cut sections containing VP were collected rostro-caudally in a series of four slides and stored at −80°C, until used for hybridization. Briefly, the day of the experiment, frozen sections were post-fixed in 4% PFA in RNA-free PBS (hereafter referred to as PBS) at room temperature (RT) for 15’, then washed in PBS, dehydrated using increasing concentration of ethanol (% in water: 50% once, 70% once, 100% twice) for 5’. Sections were then dried at RT and incubated with Protease IV for 30’ at RT. Sections were washed in PBS three times for 5’ at RT, then hybridized. Probes against *Gad2* (Cat. No. #439371, dilution 1:50) and *Slc16a7* (*Vglut2*) (Cat. No. #319171-C3, dilution 1:50) were applied to VP sections. Hybridization was carried out for 2h at 40°C. After that, sections were washed twice in PBS at RT for 2’, then incubated with three consecutive rounds of amplification reagents (30’, 15’ and 30’, respectively, at 40°C). After each amplification step, sections were washed twice in PBS at RT for 2’, twice. Finally, fluorescence detection was carried out for 15’ at 40°C. Fluorescent dyes used were Alexa-488 (for *Gad2* detection) and Atto-647 (for *Slc17a6* detection). CTB555 signal was detected in the red channel.

Sections were then washed twice in PBS, incubated with DAPI (1:10000 in PBS) for 2’, washed twice in PBS for 2’, then mounted with cover-slip using mounting medium. Images were acquired using an LSM780 confocal microscope and 20x or 40x lenses, and visualized and processed using ImageJ and Adobe Illustrator.

### Monosynaptic tracing with pseudotyped rabies virus

Retrograde tracing of monosynaptic inputs onto genetically-defined cell populations of the VP was accomplished using a previously described method (Callaway and Luo, 2015; Penzo et al., 2015). In brief, *Vglut2-Cre;Rosa26-stop^flox^-tTA* or *GAD2-Cre;Rosa26-stop^flox^-tTA* mice that express tTA in glutamatergic or GABAergic cells, respectively, were injected into the VP with AAV-TRE-hGFP-TVA-G (0.2–0.3 µl) that expresses the following components in a tTA-dependent manner: a fluorescent reporter histone GFP (hGFP); TVA (which is a receptor for the avian virus envelope protein EnvA); and the rabies envelope glycoprotein (G). Two weeks later, mice were injected in the same VP location with the rabies-EnvA-SAD-DG-mCherry (500 nl), a rabies virus that is pseudotyped with EnvA, lacks the envelope glycoprotein, and expresses mCherry. This method ensures that the rabies virus exclusively infects cells expressing TVA. Furthermore, complementation of the modified rabies virus with envelope glycoprotein in the TVA-expressing cells allows the generation of infectious particles, which then can trans-synaptically infect presynaptic neurons.

### Classical conditioning task

Four *GAD2-Cre;Ai40D* and two *Vglut2-Cre;Ai40D* mice were trained on an auditory classical conditioning task. One week after surgery mice were water-deprived in their home-cage. During training, mice were head restrained using custom-made clamps and metal head-bars. Each mouse was habituated to head restraint for one day prior to training. There were five possible outcomes (unconditioned stimuli, US), each associated with a different auditory cue (conditioned stimulus, CS): a large water reward (5 µl), a small water reward (1 µl), nothing, a small air puff (100 ms) or a large air puff (500 ms). The air puff was delivered to the animal’s face. Each trial began with a houselight turning on (the light stayed on for 5 seconds). A CS (1 second sound) was played one second after the houselight was turned on, followed by a 0.5 second delay and then a US (the outcome). In each session, reward and punishment trials were presented in two sequential blocks, with each cue chosen pseudorandomly. Each block contained the neutral stimulus. We defined satiated trials as trials where the mouse licked two or less times in the choice window.

For recording in the modified ‘conflict task’ (see below) the same auditory classical conditioning paradigm was used, except that one CS predicted a water reward (12 µl), another CS predicted the simultaneous delivery of both the water reward and a 100-ms air puff blowing to the face (the trials with different CSs were randomly interleaved).

### Reward-and-punishment conflict task

Mice were first trained and tested in the reward-only task, and subsequently trained and tested in the reward-and-punishment conflict task (or ‘conflict task’). In each trial of the reward-only task, one of five distinct auditory cues (CS; 1-s duration, presented pseudo-randomly) was presented, followed by a 500-ms delay and then an outcome (US). Each CS uniquely predicted one of five sizes of water reward: 3, 5, 7, 10 or 12 µl. Water delivery was contingent on mice licking the waterspout during a 1-s choice window, which spanned the last 500 ms of CS and the 500-ms delay after CS ended. Failure of licking during the choice window led to omission of water reward. Mice were trained for 4-8 weeks with one session per day (250 trials per session; inter-trial-interval, 8 s) until they achieved reliable licking during the choice window in trials predicting large reward. In the optogenetic testing day, animals were subjected to 250 trials, wherein laser stimulation occurred in 20% of randomly interleaved trials (laser stimulation started from CS onset and lasted until the time of water delivery). Next, mice were trained in the ‘conflict task’, which was similar to the reward-only task except that licking the waterspout during the choice window triggered delivery of both the water reward and a 100-ms air puff blowing to the face. Mice were trained for 1-2 weeks in the conflict task prior to the optogenetic testing.

### Run-for-water task

Mice were water deprived for a day prior to training in the run-for-water (RFW) task. After being habituated to the head-fixation apparatus and the running wheel, mice were presented with a CS (12-kHz, 1-s tone) that predicted the conditional availability of water (10 µl) in a spout close to the mouth. Only if the mice reached a running velocity of 10 cm/s in a response window spanning from tone onset to 500 ms after the tone ended. Failure to reach the velocity threshold during the response window resulted in water omission. Animals were trained in one session per day for 4-8 weeks (100 trials per session; average inter-trial-interval, 30 s) until they reached a reliable running response to the CS (2 z-scores above baseline running) before optogenetic testing. For the optogenetic testing, animals received 100 trials, where 20 randomly interleaved trials included laser delivery that covered the period from CS onset to the onset of water delivery. Throughout the training and testing, animals were ensured to receive a total of 1 ml of water every day.

### Run-to-avoid-air-puff task

Mice were habituated to the head-fixation apparatus and the running wheel prior to running-to-avoid-air-puff task (RTAA) training. Mice were then presented with auditory tones (white noise, 1 s) that predicted delivery of an air puff to the face (400 ms) if the animal did not reach a running speed of 10 cm/s at some point from tone onset to 500 ms after tone offset. Failure to reach the speed threshold resulted in punishment delivery. The intertrial interval was 30 s on average. Animals were trained every day for one session of 100 trials. Animals were trained from 4-8 weeks until they exhibited a reliable running response to the tone (2 z-scores above baseline running) before optogenetic testing. For the optogenetic testing day, animals received 100 trials, where 20 randomly interleaved trials included laser delivery that covered from tone onset to the air puff delivery time.

### Real-time place preference or aversion task

Freely moving mice were initially habituated to a two-sided chamber, and were subsequently subjected to a 10-min session in which laser stimulation (laser power, 10 mW (measured at fiber tip); laser frequency, 40 Hz for ChR2 experiments or the laser was constantly on for ArchT experiments) was turned on once mice entered the left side of the chamber. This was followed by another 10-min session in which laser simulation was turned on once mice entered the right side of the chamber. We recorded the percentage of time the mice spent on either side of the chamber during baseline and during the laser stimulation.

### Self-stimulation

Freely moving mice were placed in a chamber equipped with two nose-poke ports. Nose-poking to one of the ports (the active port) triggered laser delivery (duration, 2 s, intensity, 10 mW (measured at fiber tip), frequency, 40 Hz), while nose-poking to the other port (the inactive port) did not trigger laser delivery. Mice were allowed to freely poke the two ports and were tested in a single 1-hr session.

### *In vivo* electrophysiology

Custom-built screw-driven microdrives with 4 implantable tetrodes and a 50 µm fibre-optic were used to record simultaneously from multiple neurons. Each tetrode was glued to the fibre-optic with epoxy, such that the end of each tetrode was 200-400 µm from the end of the fibre-optic. Neural recordings and time stamps for behavioural variables were acquired with a Tucker-Davis Technologies RZ recording system (with a 32 channel preamp PZ2-32 and a RZ5D Bioamp processor; Alachua, Florida, USA).

Broadband signals from each wire were filtered between 0.2 and 8,500 Hz and recorded continuously at 25 kHz. To extract the timing of spikes, signals were band-pass-filtered between 300-5,000 Hz. Data analyses were carried out using software in Matlab (The Mathworks, Inc., Natick, Massachusetts, USA). Spike waveforms were manually sorted offline based on amplitude and waveform energy features using MClust-3.5 (from Dr. A. David Redish, University of Minnesota, Minneapolis, Minnesota, USA). Individual neurons were only included in the dataset if they were well isolated based on their isolation distance (>20) and L-ratio (<0.1) (Schmitzer-Torbert et al., 2005). Prior to implantation, tetrodes were dipped in DiI to aid the post-hoc visualization of the recording locations.

In order to convert raster plots of firing rate into continuous spike density functions, spike times were first binned into 1 ms time windows and then convolved with a Gaussian kernel (σ = 15 ms). To determine the response to the CS or US presentation, the average firing rates were calculated using a 300 ms window defined as 180-480 ms following the stimulus. These time windows were chosen to cover the time of the peak neuronal response. Average baseline firing was calculated using a 300 ms window immediately preceding the delivery of the CS.

To identify putative GABAergic or glutamatergic VP neurons, we used ArchT-mediated optic tagging (Courtin et al., 2014), whereby 200 ms light pulses (λ = 532 nm; OEM Laser Systems Inc., Bluffdale, Utah, USA) were delivered every 5 seconds for 100 trials following each behavioural recording session. In early sessions we also used 500 ms (*n* = 3) or 1 second (*n* = 1) light pulses, which tagged VP neurons in a similar way to that of the 200 ms light pulses.

In addition to their response to light, putative VP neurons were identified based on their firing pattern through a previously described unsupervised clustering approach (Cohen et al., 2012). Briefly, the first three principal components (PCs) of the Z scored firing rates of all neurons in the reward and punishment blocks were calculated using principal component analysis (PCA), with the singular value decomposition algorithm. Hierarchical clustering of the first three PCs was then performed using a Euclidean distance metric and a complete agglomeration method.

Cross-correlations between spike waveforms across sessions were used to determine whether the same unit was recorded over multiple sessions. The cross-correlations were calculated after aligning the negative peak of each waveform, averaging separately, and aligning the peaks of the averages. A conservative session-to-session cross-correlation coefficient of >0.95 was used to positively classify two sets of waveforms as belonging to the same unit. The correlation was calculated using the full duration of the spike in a window 10 ms prior to and 40 ms after the peak negative response.

CS–US indices were calculated as (CS - US)/(CS + US), where CS is the difference between the peak firing rate (maximum value of the PSTH) in the 500 ms after CS onset and the baseline firing rate, and US is the difference between the peak firing rate in the 500 ms after US onset and the baseline firing rate. The baseline firing rate was calculated as the mean of the PSTH in the 0.5 s before CS onset.

To calculate receiver-operating characteristic (ROC) curves, the distributions of firing rates were compared between 1 second of activity prior to the CS presentation (baseline activity) and 1 second of activity after the CS presentation, or between the distributions of firing rates following two different cues.

### *In vitro* electrophysiology

Patch clamp recording was performed as previously described (Penzo et al., 2015). Briefly, mice were anesthetized with isoflurane before they were decapitated; their brains were then dissected out and placed in ice chilled dissection buffer (110 mM choline chloride, 25 mM NaHCO_3_, 1.25 mM NaH_2_PO_4_, 2.5 mM KCl, 0.5 mM CaCl_2_, 7.0 mM MgCl_2_, 25.0 mM glucose, 11.6 mM ascorbic acid and 3.1 mM pyruvic acid, gassed with 95% O_2_ and 5% CO_2_). An HM650 Vibrating-blade Microtome (Thermo Fisher Scientific) was then used to cut 300 μm thick coronal sections that contained the LHb. These slices were subsequently transferred to a storage chamber that contained oxygenated artificial cerebrospinal fluid (ACSF) (118 mM NaCl, 2.5 mM KCl, 26.2 mM NaHCO_3_, 1 mM NaH_2_PO_4_, 20 mM glucose, 2 mM MgCl_2_ and 2 mM CaCl_2_, at 34 °C, pH 7.4, gassed with 95% O_2_ and 5% CO_2_). Following 40 min of recovery time, slices were transferred to RT (20-24 °C), where they were continuously bathed in the ACSF.

Whole-cell patch clamp recording from LHb neurons was obtained with Multiclamp 700B amplifiers and pCLAMP 10 software (Molecular Devices, Sunnyvale, California, USA), and was visually guided using an Olympus BX51 microscope equipped with both transmitted and epifluorescence light sources (Olympus Corporation, Shinjuku, Tokyo, Japan). DR-projecting LHb neurons retrogradely labeled with CTB555 (red fluorescent) were identified and patched. The external solution was ACSF. The internal solution contained 115 mM caesium methanesulphonate, 20 mM CsCl, 10 mM HEPES, 2.5 mM MgCl_2_, 4 mM Na_2_ATP, 0.4 mM Na_3_GTP, 10 mM sodium phosphocreatine and 0.6 mM EGTA (pH 7.2).

As the acute slices were prepared from mice in which GABAergic or glutamatergic VP neurons were infected with AAV expressing ChR2-YFP, to evoke VP synaptic transmission onto LHb neurons, a blue light was used to stimulate ChR2-expressing axons originating from the VP. The light source was a single-wavelength LED system (λ = 470 nm; http://www.coolled.com/) connected to the epifluorescence port of the Olympus BX51 microscope. A light pulse of 1 ms, triggered by a TTL signal from the Clampex software, was delivered every 10 seconds to drive synaptic responses. Inhibitory or excitatory postsynaptic currents (IPSCs or EPSCs, respectively) were low-pass filtered at 1 KHz and recorded. IPSCs were recorded at a holding potential of 0 mV, with APV (100 μM) and CNQX (5 μM) added to the ACSF. EPSCs were recorded at holding potentials of –70 mV (for AMPA-receptor-mediated responses) and +40 mV (for NMDA-receptor-mediated responses), with picrotoxin (100 μM) added to the ACSF. Synaptic responses were analyzed using pCLAMP 10 software.

### Statistics and data presentation

To determine whether parametric tests could be used, the Shapiro-Wilk Test was performed on all data as a test for normality. The statistical test used for each comparison is indicated when used. The sample sizes used in this study were based on estimations by a power analysis. Behavioural tests and electrophysiological data acquisition were performed by investigators with knowledge of the identities of experimental groups. All these experiments were controlled by computer systems, with data collected and analysed in an automated and unbiased way. No data points were excluded.

## SUPPLEMENTARY FIGURES

**Supplementary Figure 1.**
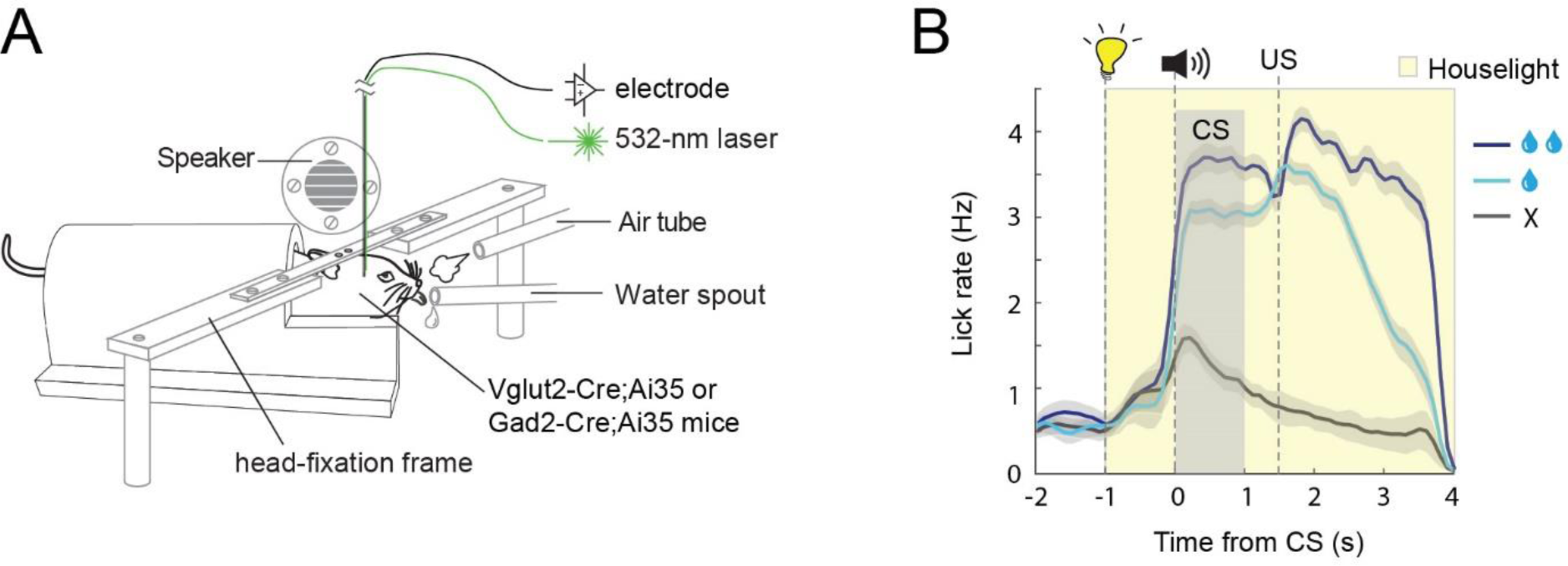
The behavioural task for in vivo recording. **(A)** A schematic of the experimental design. **(B)** Average licking behaviour in the reward conditioning task (n=14 sessions from 4 mice), P < 0.01 between large and small reward trials, P < 0.001 between small and no reward trials (paired T-test). Shaded areas indicate s.e.m.

**Supplementary Figure 2.**
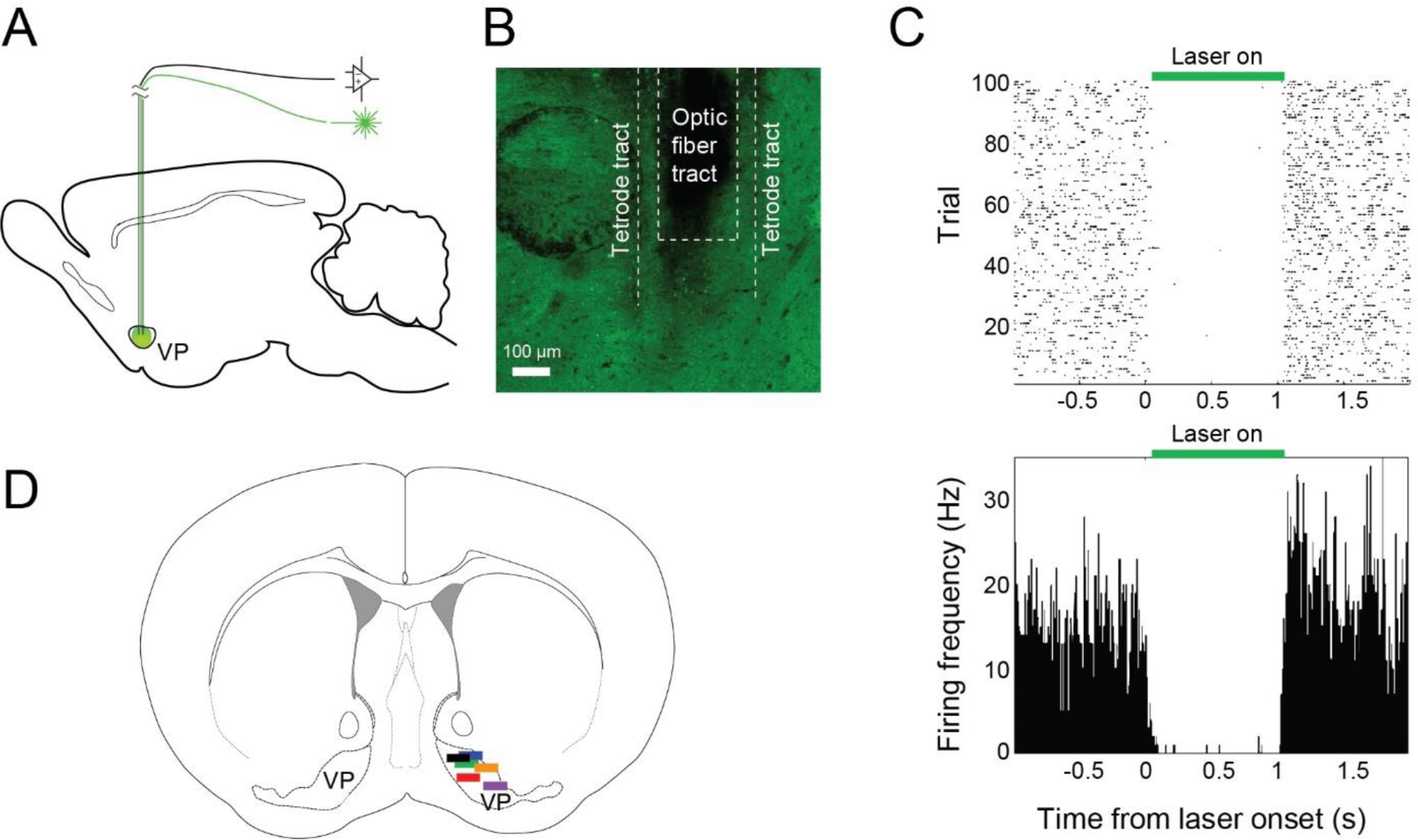
In vivo recording and optogenetic tagging. **(A)** A schematic of the experimental approach. **(B)** Optic fibre and tetrode tract showing the recording location in the VP in a representative mouse. **(C)** Raster plot (top) and peri-stimulus time histogram (bottom) of an example tagged neuron. **(D)** A schematic showing the recording locations in 6 mice.

**Supplementary Figure 3.**
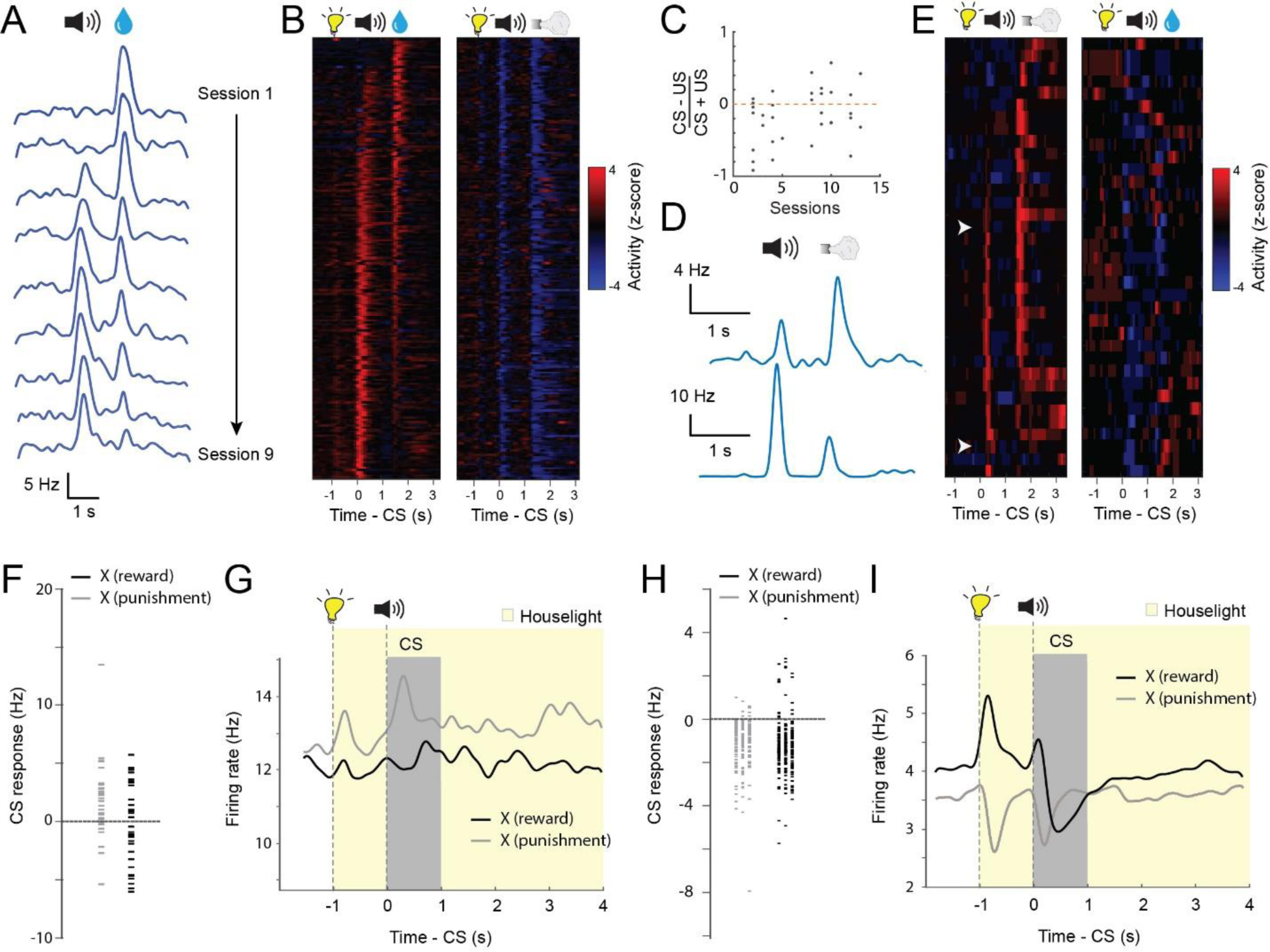
Development of responses in VP neurons during learning. **(A)** The responses to reward cue (CS) or reward (US) in an example PVN tracked over multiple sessions (S1-S9). Responses are shown as spike density plots. **(B)** Z-score activity plots of the responses of all PVNs during reward and punishment blocks. Each row represents the activities of one neuron. Neurons are sorted according to their CS/US response ratio. **(C)** The CS-US response index for all NVNs in the punishment block across different stages of training (r^2^ = 0.41, P < 0.05 by linear regression). **(D)** The CS and US responses of two example NVNs in the punishment block at different stages of training. Responses are shown as spike density plots. **(E)** Z-score activity plots of the responses of all NVNs during reward and punishment blocks. Each row represents the activities of one neuron. Neurons are sorted according to their CS/US response ratio. **(F)** Graph showing the CS responses of all NVNs during the neutral cue trials in the reward and punishment blocks (reward block, mean, −0.32 Hz; punishment block, mean, −1.96 Hz). **(G)** Average responses of NVNs during neutral trials in reward and punishments blocks. Responses are shown as spike density plots. **(H)** Graph showing the CS responses of all PVNs during the neutral cue trials in the reward and punishment blocks (reward block, mean, −1.26 Hz; punishment block, mean, −1.05 Hz) **(I)** Average responses of PVNs during neutral trials in reward and punishments blocks. Responses are shown as spike density plots.

**Supplementary Figure 4.**
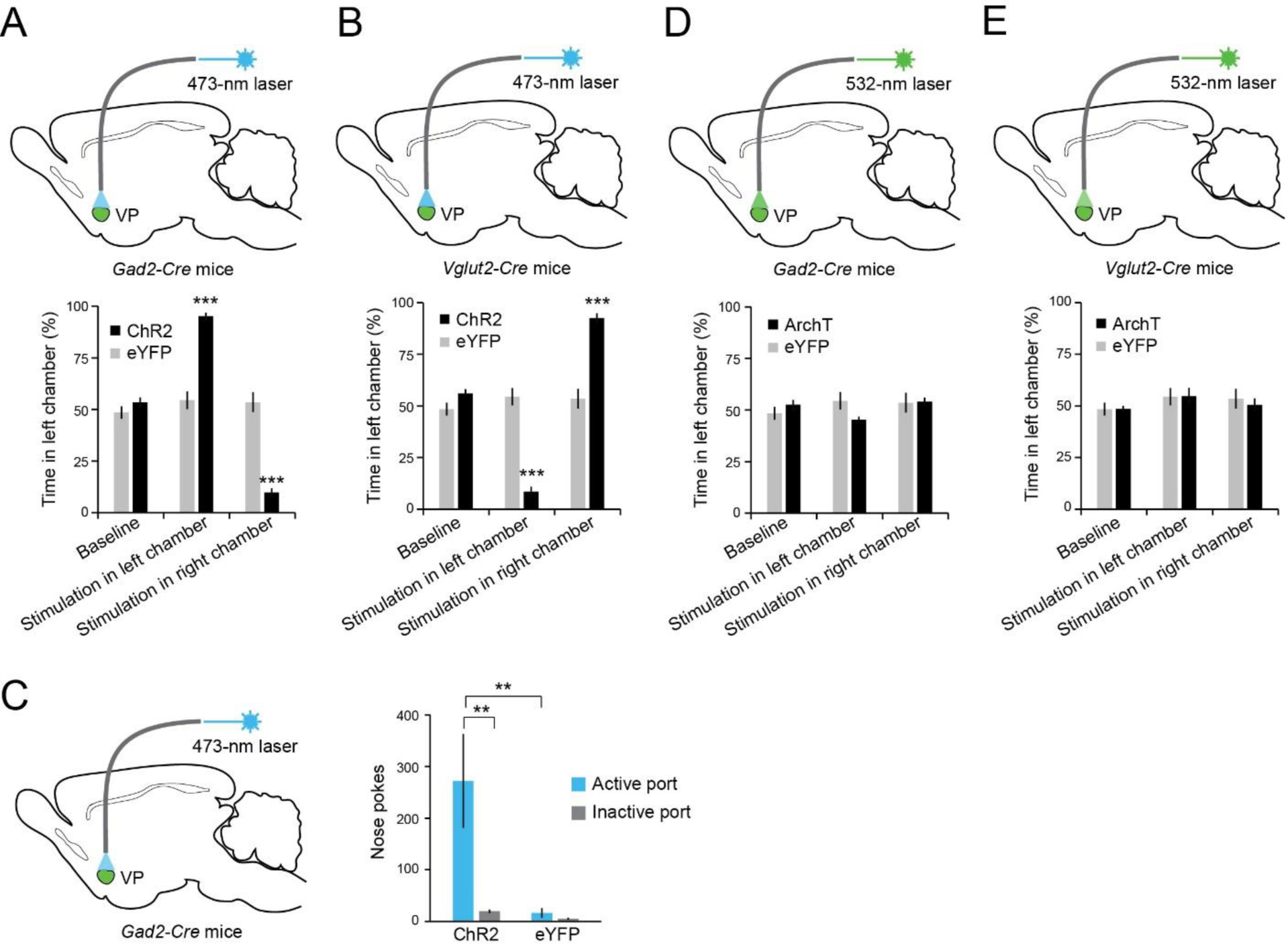
Activation of GABAergic or glutamatergic VP neurons is intrinsically rewarding or aversive, respectively. **(A)** Top: a schematic of the approach. Bottom: activation of GABAergic VP neurons induced real-time place preference (F_(5, 26)_ = 81.86, p < 0.0001, n = 4, 5). ***P < 0.001, two-way ANOVA followed by Bonferroni’s test. **(B)** Top: a schematic of the approach. Bottom: activation of glutamatergic VP neurons induced real-time place aversion (F_(5, 26)_ = 59.56, p < 0.0001, n = 4, 5). ***P < 0.001, two-way ANOVA followed by Bonferroni’s test. **(C)** Left: a schematic of the approach. Right: activation of GABAergic VP neurons supported self-stimulation (F_(3, 15)_ = 8.71, p = 0.012, n = 4, 4). **P < 0.01, two-way ANOVA followed by Tukey’s test. **(D)** Top: a schematic of the approach. Bottom: inactivation of GABAergic VP neurons had no effect on place preference (F_(5, 29)_ = 0.63, p = 0.54, n = 5, 5, two-way ANOVA). **(E)** Top: a schematic of the approach. Bottom: inactivation of glutamatergic VP neurons had no effect on place preference (F_(5, 29)_ = 1.39, p = 0.27, n = 5, 5, two-way ANOVA). Data are presented as mean ± s.e.m.

**Supplementary figure 5.**
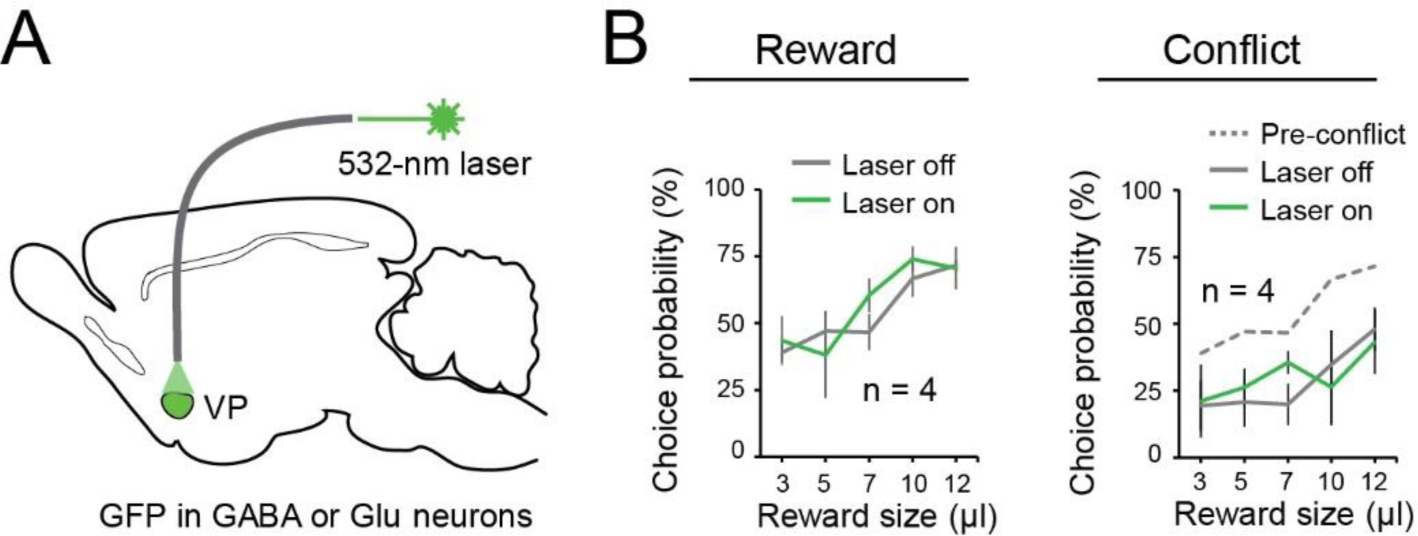
VP GFP control mice in the reward and conflict tasks. **(A)** A schematic of the approach. **(B)** Laser delivery into the VP had no effect on reward seeking in the reward task (F_(1, 7)_ = 0.39, p = 0.54) (left), or in the conflict task (F_(1, 7)_ = 0.17, p = 0.68) (right) in these GFP controls. Two-way ANOVA. Data are presented as mean ± s.e.m.

**Supplementary figure 6.**
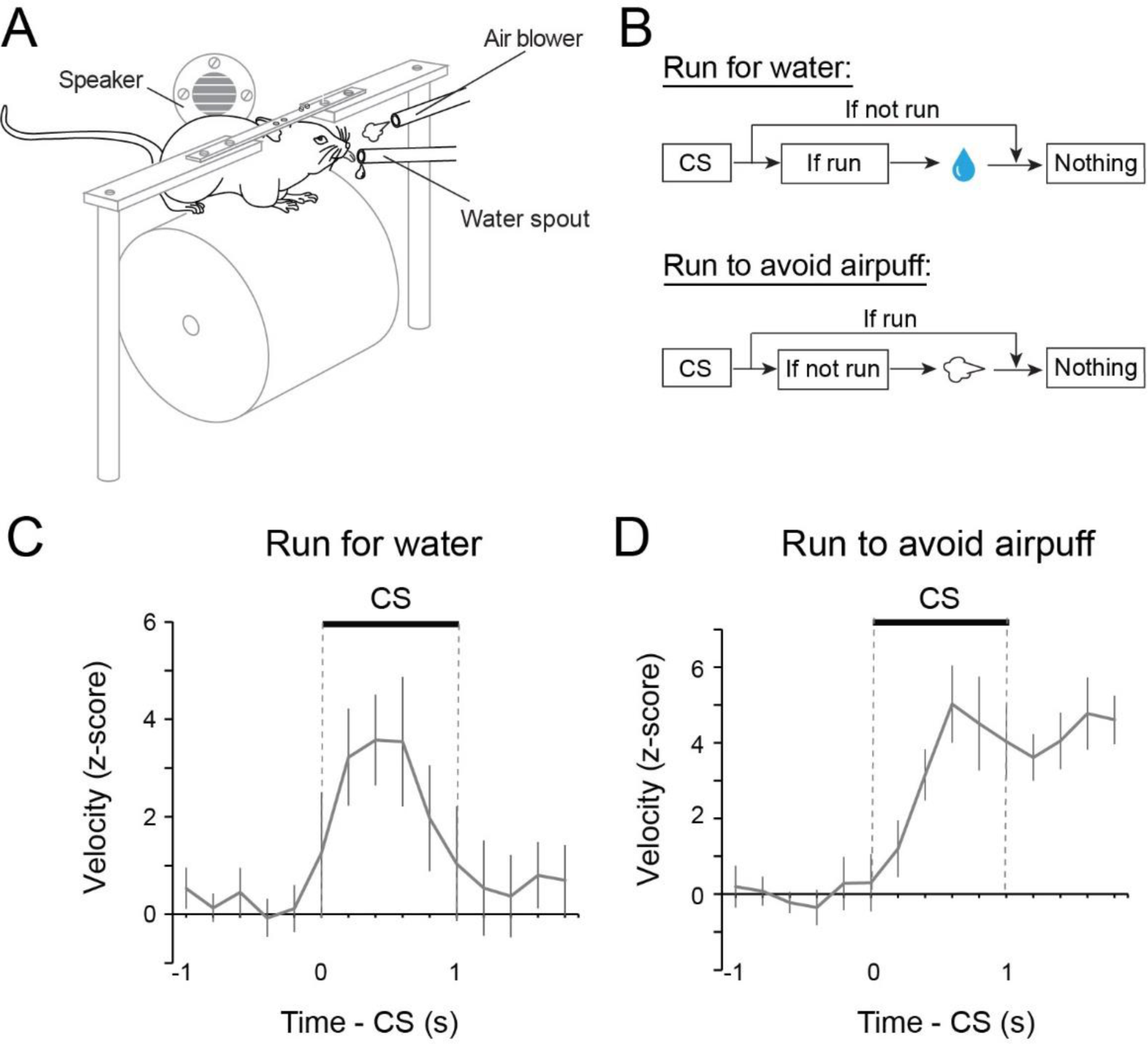
The running tasks. **(A)** A schematic of the experimental design. **(B)** Schematics of the experimental procedure. **(C)** Behavioral performance of mice in the run-for-water task (n = 7). **(D)** Behavioral performance of mice in the run-to-avoid-air-puff task (n = 4). Data are presented as mean ± s.e.m.

**Supplementary figure 7.**
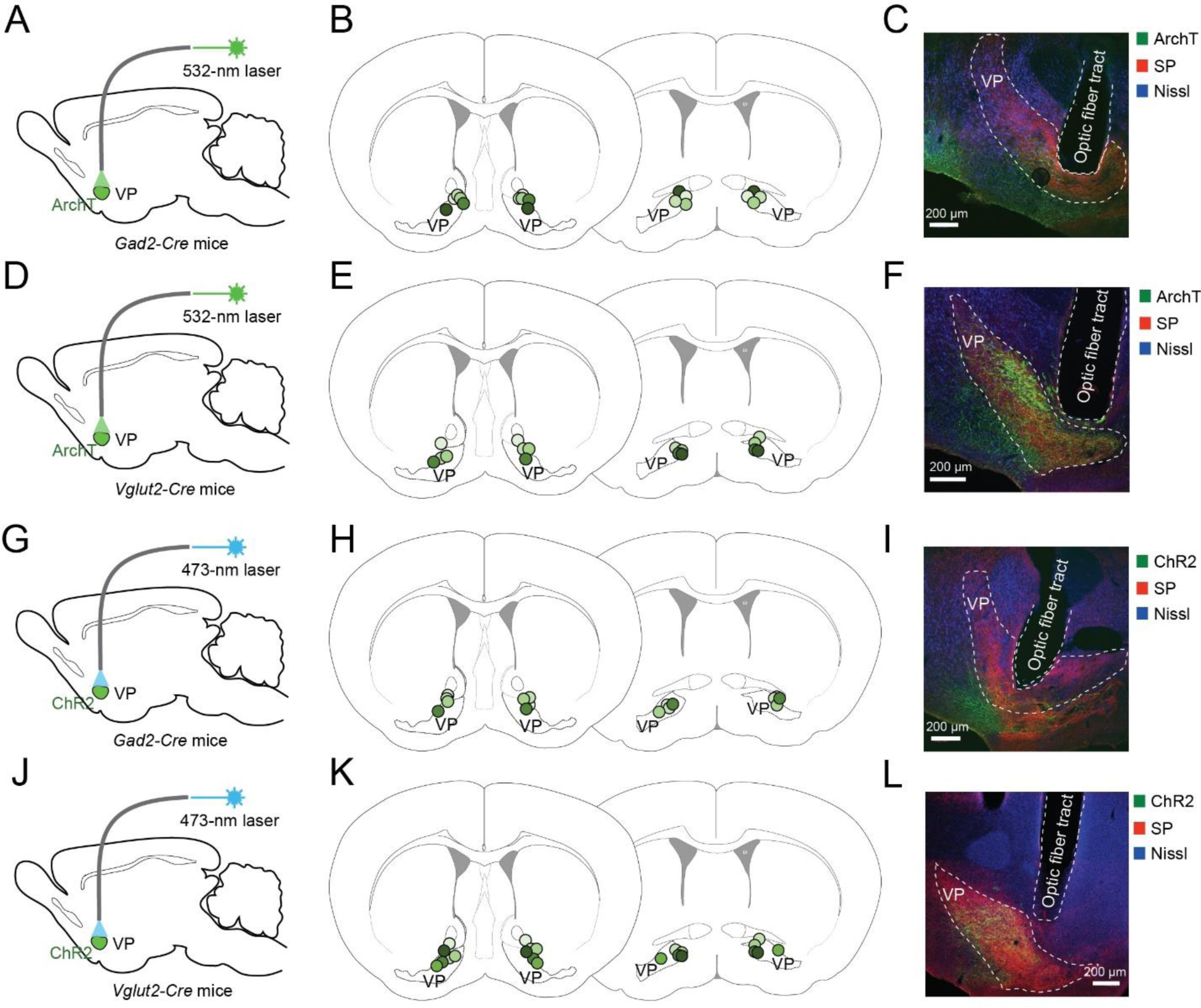
Histology results for mice in which optogenetic manipulation was performed in the VP. **(A-F)** Histology results for mice in which VP GABAergic (A-C) or glutamatergic (D-F) neurons were optogenetically inhibited. Left: schematics of the approach. Middle: schematics of optical fibre placement locations. Right: images of the VP from representative mice, showing the ArchT-GFP-expressing VP neurons and the expression of Substance P (SP), which was used as a marker to delineate the VP. **(G-L)** Histology results for mice in which VP GABAergic (G-I) or glutamatergic (J-L) neurons were optogenetically activated. Left: schematics of the approach. Middle: schematics of optical fibre placement locations. Right: images of the VP from representative mice, showing the ChR2-eYFP-expressing VP neurons and the expression of SP.

**Supplementary figure 8.**
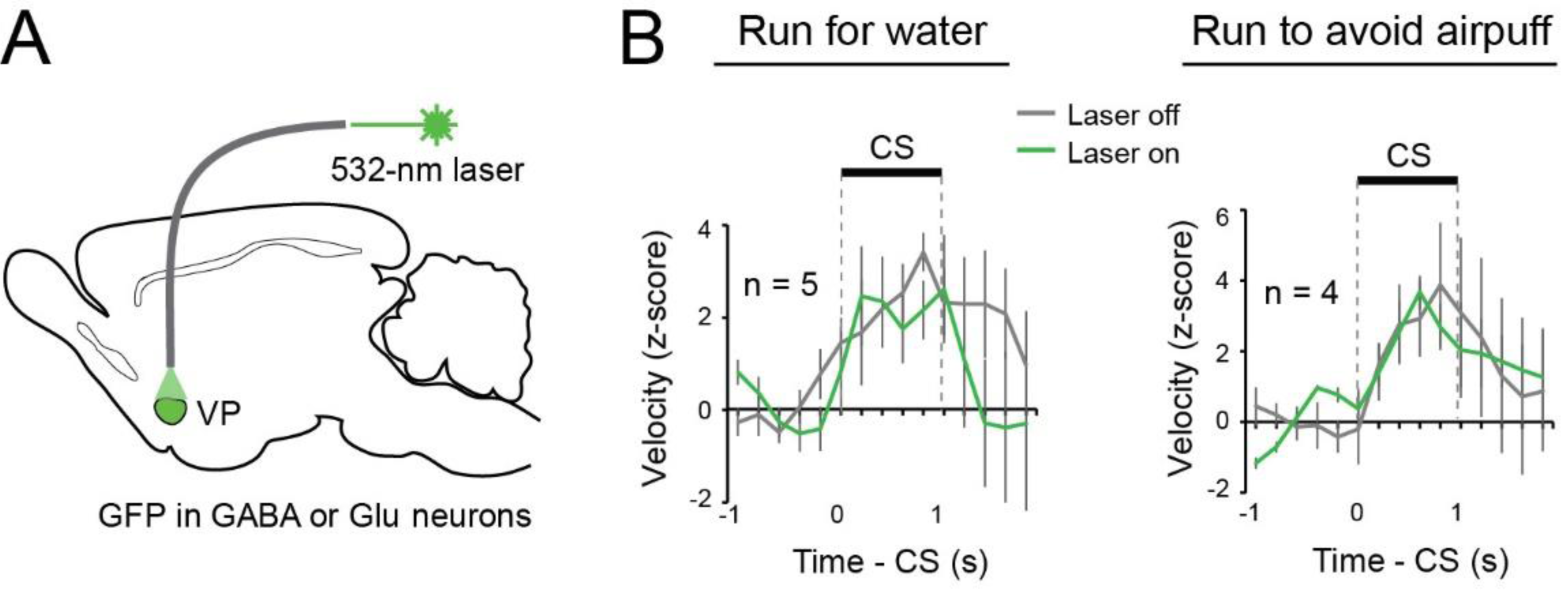
VP GFP control mice in the running tasks. **(A)** A schematic of the approach. **(B)** Laser delivery into the VP had no effect on running for water (F _(1, 9)_ = 3.17, p = 0.080) (left), or on running to avoid air puff (F_(1, 7)_ = 0.0016, p = 0.97) (right) in these GFP controls. Two-way ANOVA. Data are presented as mean ± s.e.m.

**Supplementary figure 9.**
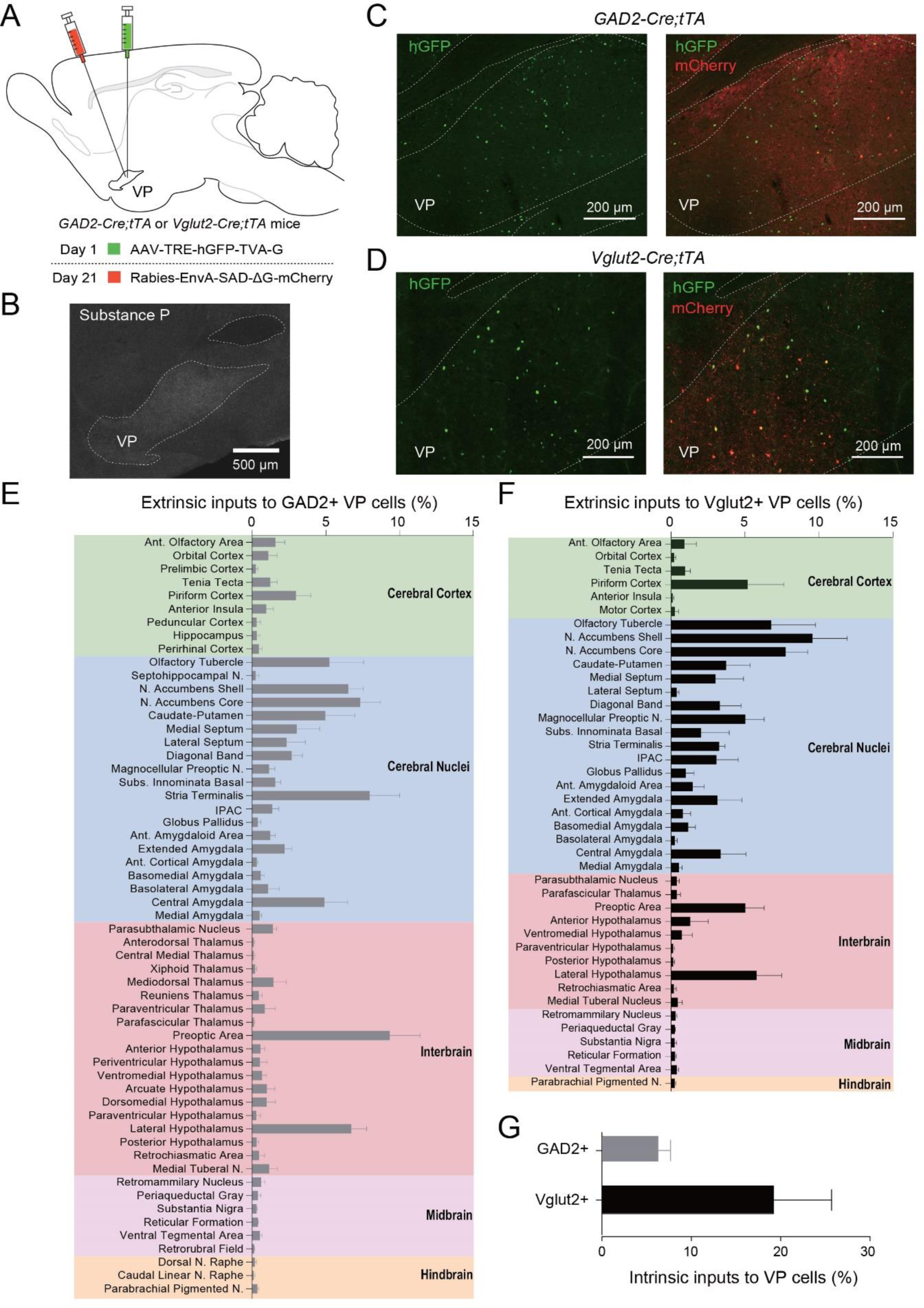
Monosynaptic inputs onto GABAergic and glutamatergic VP neurons. **(A)** Schematics of the experimental design. GABAergic or glutamatergic VP neurons were targeted using GAD2-Cre;Rosa26-stopflox-tTA or Vglut2-Cre;Rosa26-stopflox-tTA mice, respectively. **(B)** The expression of Substance P (SP), recognized by an antibody, was used to identify the VP. **(C, D)** Confocal images showing the GABAergic **(C)** and glutamatergic **(D)** starter cell (yellow) location in the VP. **(E, F)** Graphs showing the fraction of monosynaptically labelled neurons in each brain region that projected to the GABAergic **(E)** (n = 4 mice) or glutamatergic **(F)** (n = 4 mice) VP neurons. **(G)** The fraction of local VP neurons that provided monosynaptic inputs onto VP GABAergic or glutamatergic neurons. Data in E-G are presented as mean ± s.e.m.

**Supplementary figure 10.**
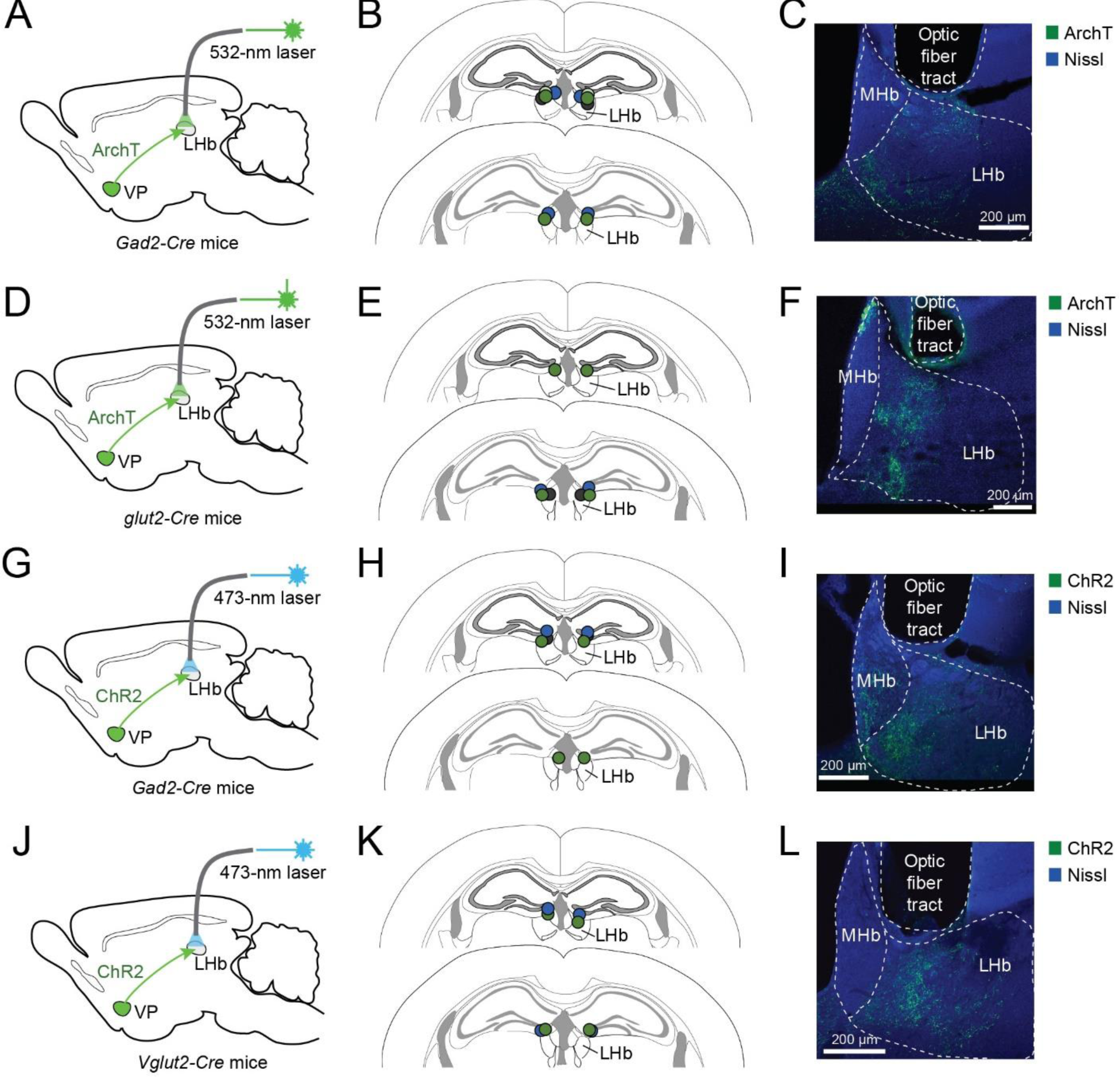
Histology results for mice in which optogenetic manipulation was performed in the VP-LHb pathway. **(A-F)** Histology results for mice in which the GABAergic^VP→LHb^ (A-C) or glutamatergic^VP→LHb^ (D-F) pathways were optogenetically inhibited. Left: schematics of the approach. Middle: schematics of optical fibre placement locations. Right: images of the LHb from representative mice, showing the ArchT-GFP-expressing axons originating from the VP. **(G-L)** Histology results for mice in which the GABAergic^VP→LHb^ (G-I) or glutamatergic^VP→LHb^ (J-L) pathways were optogenetically activated. Left: schematics of the approach. Middle: schematics of optical fibre placement locations. Right: images of the LHb from representative mice, showing the ChR2-eYFP-expressing axons originating from the VP.

**Supplementary figure 11.**
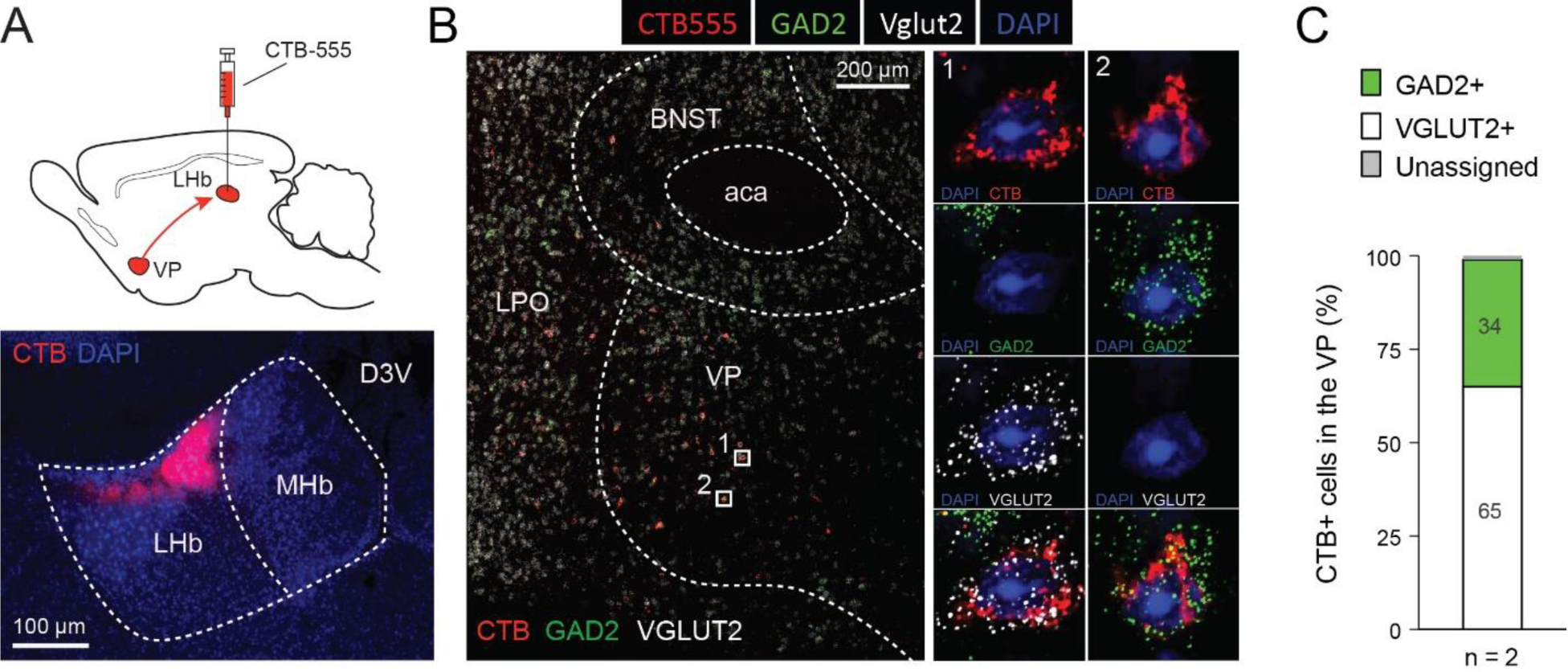
Both GABAergic and glutamatergic VP neurons project to the LHb. **(A)** Top: a schematic of the approach. Bottom: an image of the LHb from a representative mouse, showing the location of CTB555 (red) injection site in the LHb. **(B)** Low (left) and high (right) magnification confocal images of the VP from a representative mouse, demonstrating fluorescent single molecule in situ hybridization for GAD2 (green) and Vglut2 (white), as well as DAPI staining (blue) and the retrograde labelling by CTB555 (red) from the LHb. **(C)** Quantification of VP neurons co-labeled with CTB and GAD2 or VGlut2.

**Supplementary figure 12.**
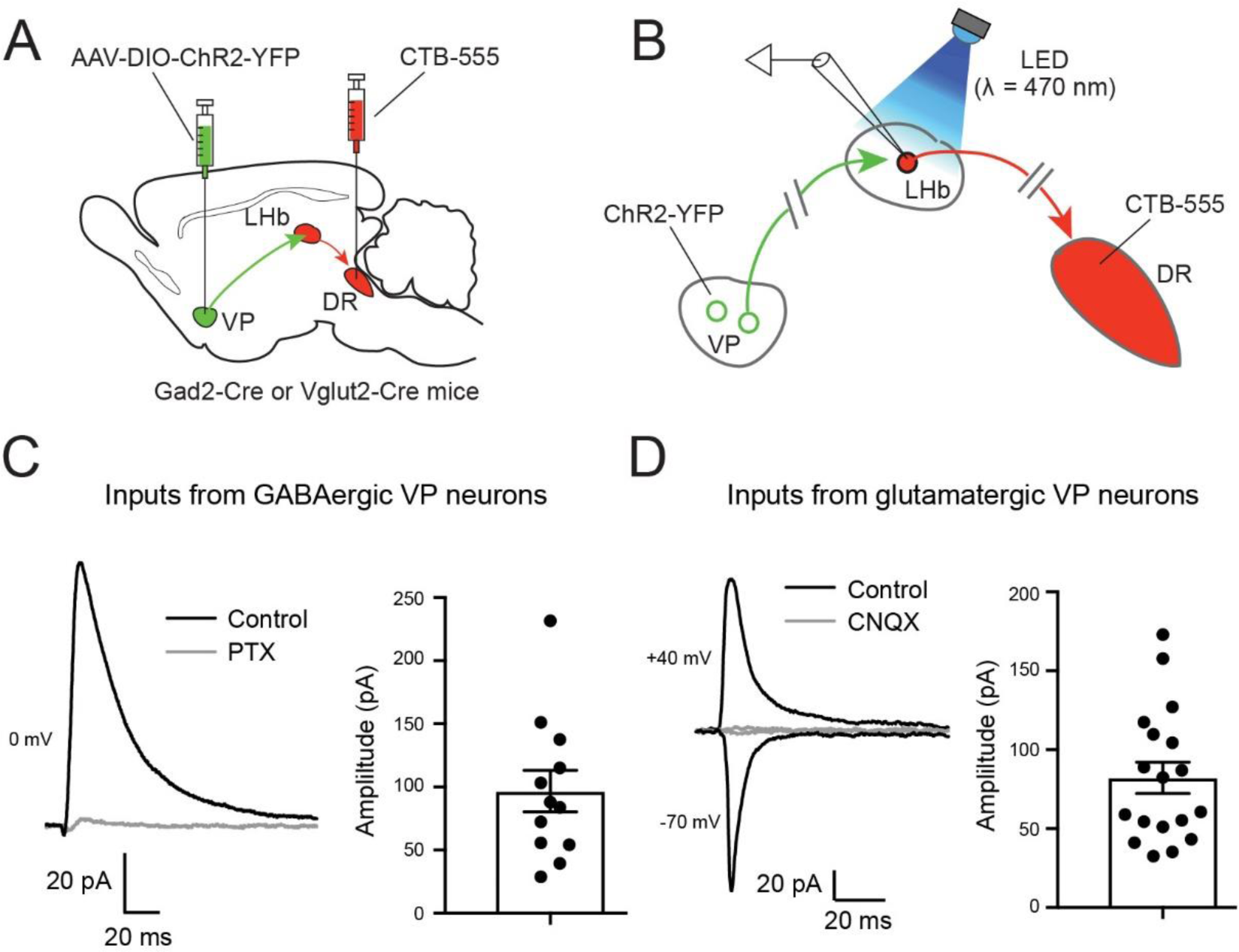
The DR-projecting LHb neurons receive both inhibitory and excitatory synaptic inputs from the VP. **(A, B)** Schematics of the experimental approach. **(C)** Left: traces of inhibitory postsynaptic currents (IPSCs) recorded from a DR-projecting LHb neurons with whole-cell patch clamp recording. Right: quantification of IPSC amplitude for all recorded neurons (12 cells from 2 mice). **(D)** Left: traces of excitatory postsynaptic currents (EPSCs) recorded from a DR-projecting LHb neurons with whole-cell patch clamp recording. Right: quantification of EPSC amplitude for all recorded neurons (18 cells from 3 mice). Data in C, D are presented as mean ± s.e.m.

**Supplementary Figure 13.**
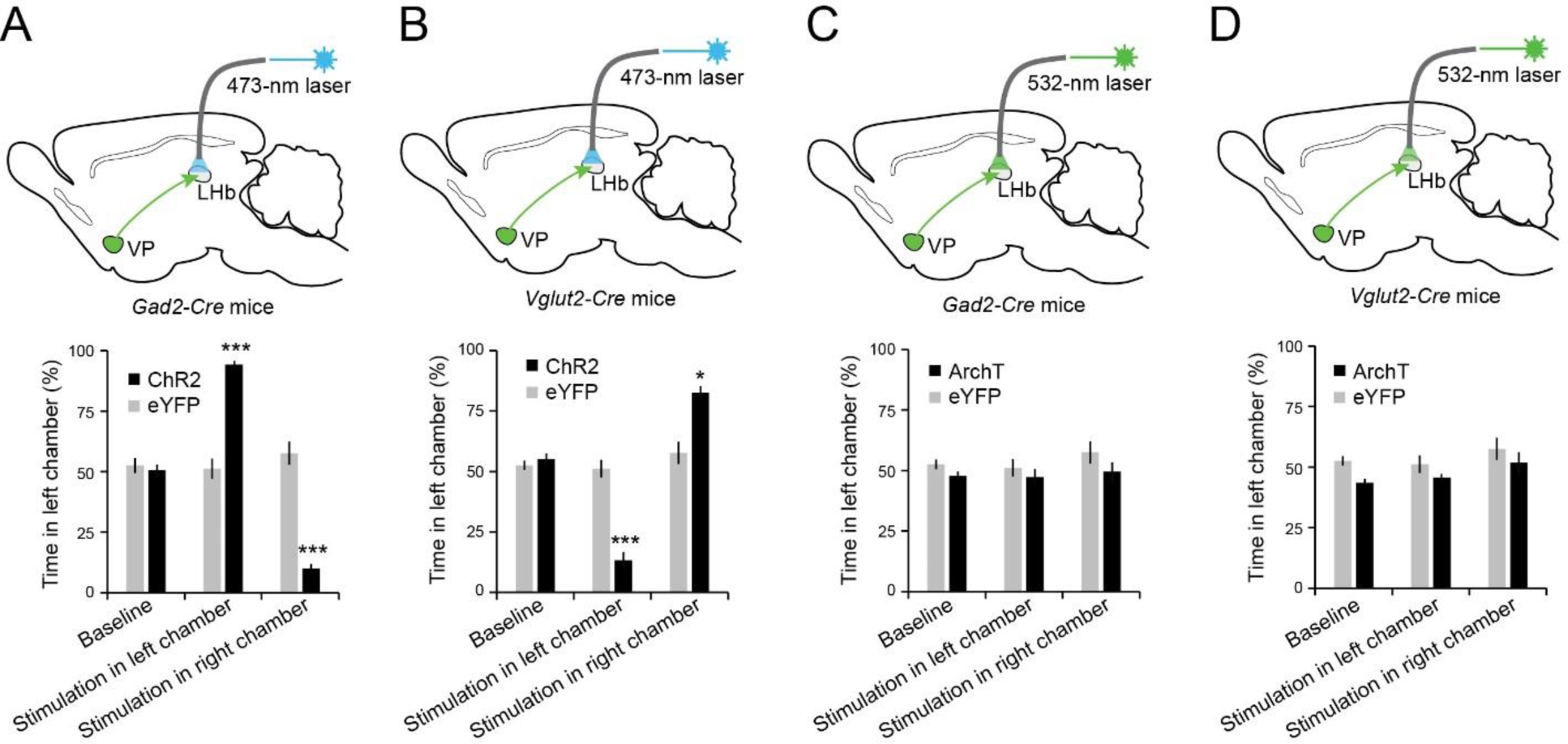
Activation of GABAergic^VP→LHb^ or glutamatergic^VP→LHb^ projections is intrinsically rewarding or aversive, respectively. **(A)** Top: A schematic of the approach. Bottom: activation of the GABAergic^VP→LHb^ projections induced real-time place preference (F_(5,29)_ = 75.74, p < 0.0001, n = 4, 6). ***P < 0.001, two-way ANOVA followed by Bonferroni’s test. **(B)** Top: A schematic of the approach. Bottom: activation of the glutamatergic^VP→LHb^ projections induce real-time place aversion (F_(5,29)_ = 86.84, p < 0.0001, n = 4, 6). *P < 0.05, ***P < 0.001, two-way ANOVA followed by Bonferroni’s test. **(C)** Top: A schematic of the approach. Bottom: inactivation of the GABAergic^VP→LHb^ projections did not induce place preference or aversion (F_(5, 35)_ = 0.93, p = 0.41, n = 6, 6). Two-way ANOVA. **(D)** Top: A schematic of the approach. Bottom: inactivation of the glutamatergic^VP→LHb^ projections did not induce place preference or aversion (F_(5, 29)_ = 1.83, p = 0.19, n = 4, 6). Two-way ANOVA. Data are presented as mean ± s.e.m.

**Supplementary figure 14.**
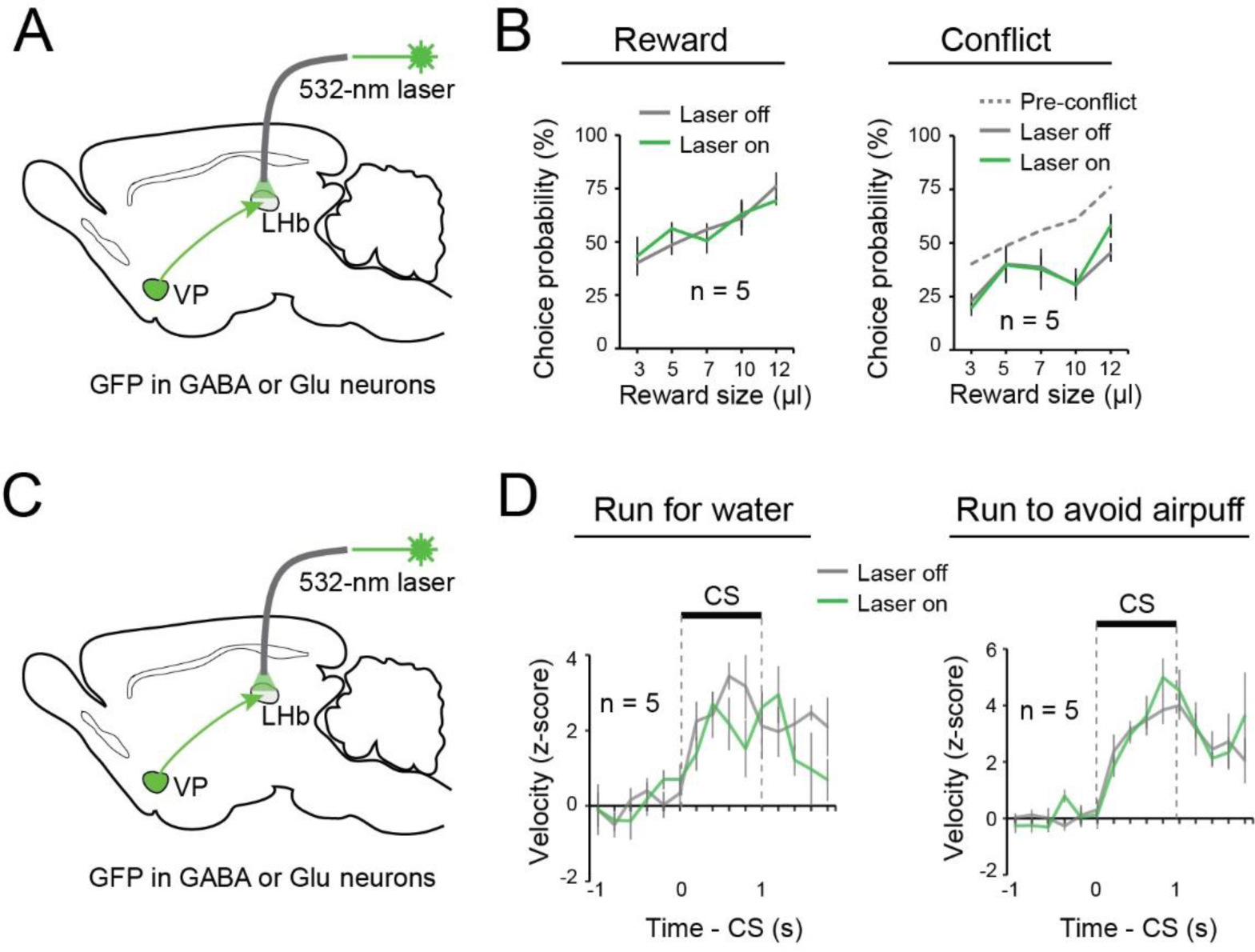
GFP control mice for the VP-LHb pathway in the reward and conflict tasks and running tasks. **(A)** A schematic of the approach. **(B)** Laser delivery into the LHb had no effect on reward seeking in the reward task (F_(1, 9)_ = 0.0015, p = 0.97) (left), or in the conflict task (F_(1, 9)_ = 0.19, p = 0.67) (right) in these GFP controls. Two-way ANOVA. **(C)** A schematic of the approach. **(D)** Laser delivery into the LHb had no effect on running for water (F_(1, 9)_ = 3.26, p = 0.074) (left), or running to avoid air puff (F_(1, 9)_ = 0.40, p = 0.53) (right) in these GFP controls. Two-way ANOVA. Data are presented as mean ± s.e.m.

